# Visualising diverse extracellular nucleic acids in biofilms with DNA-binding dyes

**DOI:** 10.64898/2026.05.08.723897

**Authors:** Freja Winther Sillesen, Finn Dicke, Stephanie Kath-Schorr, Hannah Weissinger, Jørgen Kjems, Amalia Villum Jakobsen, Daniel Erik Otzen, Gabriel Antonio Salvador Minero, Rikke Louise Meyer

## Abstract

Extracellular nucleic acids (eNA) are central components of bacterial biofilms, contributing to structural integrity, antibiotic tolerance, and emerging functions such as extracellular electron transfer and peroxidase-like catalysis. While extracellular DNA has traditionally been assumed to adopt the canonical B-DNA conformation, biofilms are now known to contain non-canonical structures, including Z-DNA/RNA (Z-NA), G-quadruplex DNA/RNA (G4-NA), and substantial amounts of extracellular RNA. Conventional nucleic acid-binding dyes are widely used for rapid eNA detection, yet their specificity for these diverse structures has not been systematically evaluated. Here, we compare the fluorescence properties of eleven membrane-impermeant dyes (TOTO™, BOBO™, YOYO™, and POPO™ series, SYTOX™ Green, SYTOX™ Red, and propidium iodide) against synthetic B-DNA, Z-DNA, G4-DNA, A-RNA, Z-RNA, and G4-RNA oligonucleotides, with Z-NA stabilised through brominated guanosine analogues synthesised in-house. A clear pattern emerged: green-fluorescent dyes preferentially bound canonical B-DNA, whereas red-fluorescent counterparts displayed broader specificity that extended to non-canonical structures. TOTO™-3 and SYTOX™ Red bound G4-NA with higher fluorescence than B-DNA, and propidium iodide showed an unexpected preference for A-RNA over B-DNA. These observations were validated in *Staphylococcus aureus* biofilms by parallel immunolabelling with structure-specific antibodies. TOTO™-3, YOYO™-3, BOBO™-3, POPO™-3, and propidium iodide reproduced the eNA distribution at the bacterial cell surface. Finally, we introduce poly-A tailing with fluorescently labelled ATP as a stringent, RNA-specific imaging method for biofilms. Together, these results provide practical guidelines for visualising the structural diversity of eNA in biofilms.

**HIGHLIGHTS:**

- Biofilms contain non-canonical structures of extracellular DNA and RNA
- This study tests the ability of DNA-binding dyes to visualise such structures
- Propidium iodide visualises RNA with brighter fluorescence than DNA
- Red-fluorescent dyes were more versatile than green-fluorescent dyes
- Combining several dyes enabled the detection of non-canonical structures

## 1. INTRODUCTION

Extracellular DNA (eDNA) has been established as a central component in biofilm formation, and it is the only matrix component reported to be shared across nearly all biofilms. In the matrix, eDNA interacts with other biopolymers, such as polysaccharides (Peng et al., 2020) and proteins (Buzzo et al., 2021), which stabilise the eDNA. Its role in bacterial biofilms has been well characterised over the last two decades, and it is critical for the biofilm’s structural integrity (Flemming & Wingender, 2010; Sharma & Rajpurohit, 2024). Some bacteria produce eDNA-degrading nucleases to combat biofilm-forming competitors in complex microbial communities (Lander et al., 2024). eDNA also impacts the physical properties of biofilms by increasing the viscoelasticity of the biofilm matrix, which allows stretching and enhances the resistance to mechanical stress (Secchi et al., 2022; Seviour et al., 2021). Recent studies have revealed new functions of eDNA in the biofilm matrix beyond structural support. These include extracellular electron transfer (EET) (Ajunwa et al., 2025; Kotoky et al., 2025; Saunders et al., 2020), peroxidase-like catalytic activity (Minero et al., 2024), and increased resistance to antibiotics (Chiang et al., 2013) and to the host immune system (Gallucci, 2024).

Traditionally, eDNA was assumed to primarily exist in the canonical right-handed double-stranded B-DNA conformation. However, recent studies have shown that biofilms form non-canonical nucleic acid structures, including left-handed double-stranded Z-DNA/RNA (Z-NA) and four-stranded G-quadruplex (G4)-DNA/RNA (G4-NA) (Buzzo et al., 2021; Evans et al., 2025; Minero et al., 2024; Seviour et al., 2021). The transition from B- to Z-NA or G4-NA often happens in guanine-rich sequences and is driven by DNA-binding proteins, high salinity, negative supercoiling, mechanical stress, and charge neutralisation, e.g., through interactions with cationic polysaccharides (Meyer et al., 2024). Furthermore, the levels of non-canonical NA structures increase as the biofilm matures (Buzzo et al., 2021). Z-NA and G4-NA are resistant to mammalian DNase I and thereby contribute to the resilience of the biofilm (Minero et al., 2024). Additionally, G4-structures can bind a plethora of small molecules, such as metalloporphyrins (e.g., heme), and acquire peroxidase-like properties in biofilms (Minero et al., 2024) and promote EET via heme bound to G4-NA. Thus, visualisation of various NA structures is essential for understanding their expanded role in biofilms.

Fluorescently labelled monoclonal antibodies (mAb) allow for bright signals, high photostability, and high specificity for Z-NA (mAb Z22 (Buzzo et al., 2021; Möller et al., 1982), mAb Z-D11 (Pohl, 1987)), G4-NA (mAb BG4 (Biffi et al., 2013; Buzzo et al., 2021; Minero et al., 2024)), B-DNA (mAb 27156 clone 35I9 (Buzzo et al., 2021; Heegaard et al., 1996; Minero et al., 2024)) and A-RNA (mAb J2 (Bou-Nader et al., 2025; Schönborn et al., 1991)) in the extracellular matrix. The cons of using mAbs are high cost and time-consuming application, and antibody penetration can be limited in thick biofilms. Furthermore, some antibodies can promote conformational changes of the DNA/RNA, causing a false-positive signal (Lee et al., 2026).

A faster method for the detection of eNA is to use membrane-impermeable nucleic acid-binding dyes that light up when bound to their target. Commonly used dyes are propidium iodide (PI) (Boulos et al., 1999; Savorana et al., 2025), SYTOX™ Green (cyanine monomer) (Roth et al., 1997; Smolarz et al., 2021), and TOTO™-1 (cyanine dimer) (Mugunthan et al., 2023; Rye et al., 1992), and a previous study evaluating the capability of different dyes to detect eDNA in biofilms recommended TOTO™-1 or SYTOX™ Green for sensitive detection (Okshevsky & Meyer, 2014).

The cyanine monomers and dimers are available in different variations with excitation and emission in green, yellow, red, and far-red ranges of the spectrum. While these DNA-binding dyes allow fast and inexpensive detection compared to immunolabelling (Lander et al., 2024), their specificity towards various eNA structures has not yet been explored. Most dyes were developed to bind to B-DNA, e.g., for use in live/dead staining, but some have been used for eDNA imaging in biofilms. With the discovery of the abundance of RNA (Chiba et al., 2022; Mugunthan et al., 2023) and non-canonical structures in the biofilm matrix, we might overlook these structures when using DNA-binding dyes.

We therefore sought to evaluate and compare the specificity of extracellular nucleic acid-binding dyes for the most common nucleic acid structures reported in biofilms: B-DNA, Z-DNA, and G4-DNA, as well as their RNA counterparts. We first assessed the dyes *in vitro* using pre-folded oligos and subsequently demonstrated their use *in situ*, in biofilms that contain Z-NA, G4-NA, and A-RNA, as confirmed by fluorescence immunolabelling. Our findings show that TOTO™-1 and SYTOX™ Green preferentially light up B-DNA, while TOTO™-3 and other red-fluorescent dyes have broader specificity towards both B-DNA and A-RNA as well as G4-NA. From the observations, we recommend using TOTO™-3 or a combination of TOTO™-3 and TOTO™-1 as a fast method for screening biofilms for eNA before identifying eNA structures with structure-specific antibodies or other structure-specific staining methods. Given the lack of DNA vs RNA specificity of dyes and even antibodies, we also present a new approach for specific detection of RNA via fluorescently labelled poly-A tailing.

## 2. MATERIALS AND METHODS

### 2.1 Nucleic acids, NA-binding dyes, and antibodies

DNA-binding dyes TOTO™-1/-3, YOYO™-1/-3, BOBO™-1/-3, POPO™-1/-3 (Dimer Sampler kit #N7565), SYTOX™ Green (#S7020), SYTOX™ Red (#S34859), and Propidium Iodide (PI, #P3566) were purchased from Thermofisher Scientific. The stock concentration of the dyes was 1 mM in DMSO and was diluted to 50 µM stocks in milliQ water, except for SYTOX™ Red and PI. SYTOX™ Red was provided as a 5 µM stock in DMSO. PI (1 mg/mL in water, *ca.* 1.5 mM) was also aliquoted as 50 µM in milliQ water. Intracellular live-dye SYTO™ 41 (Invitrogen, #S11352) was purchased from Thermo Fisher Scientific and diluted to 200 µM in milliQ water. Lipophilic dye FM 4-64 was purchased from Thermo Fisher Scientific and diluted in PBS to a concentration of 1 g/L. All aliquots were stored at -20°C.

Nucleic acid sequences are listed in Table 1.

**Table 1.** High GC-content DNA and RNA sequences used in this work to anneal various substrates for fluorescence quantification in NA-dye complexes. B-DNA, G4-DNA, A-RNA, and G4-RNA were purchased from IDT (HPLC grade). Brominated DNA and RNA to form Z-NA were synthesised on demand. C, G, T, and A stand for deoxyribonucleic cytosine, guanine, thymine, and adenine, respectively. The rC, rG, rU, and rA stand for their ribonucleic counterparts (U = uracil).

|  | Sequence | Modification |
| --- | --- | --- |
| B-DNA | 5'-C G C G C G-3' |  |
| Z-DNA | 5'-C <sup>Br</sup>G C <sup>Br</sup>G C G-3' | **BrG** = 8-Br-dG |
| G4-DNA | 5'-G A G G G T G G G T A G G G T G G G |  |
| A-RNA | 5'-rC rG rC rG rC rG-3' |  |
| Z-RNA | 5'-rC <sup>Br</sup>Gm rC <sup>Br</sup>Gm rC rG-3' | **BrG<sub>m</sub>** = 8-Br-2'-OMe-rG |
| G4-RNA | 5'-rG rA rG rG rG rU rG rG rG rU rA rG rG rG rT rG rG rG |  |

Monoclonal antibodies BG4 (goat, T2206D03) and Z22 (rabbit, T2307A04) were purchased from Vector Labs (former Absolute Antibodies), conjugated with Atto488 or FluoroProbes647h. Monoclonal anti-B-DNA antibody (“AB1” clone 35I9, mouse, 1123857-2) was purchased from Abcam. Monoclonal anti-A-RNA antibody (J2, *RNT-SCI-10010200*) was purchased from Jena Bioscience.

### 2.2 Synthesis of brominated Z-DNA and RNA

For in-house solid-support oligonucleotide synthesis, an H-6 DNA/RNA synthesiser from K&A Labs GmbH was used. As solid support for DNA synthesis, preloaded dG-CPG was purchased from LGC Biosearch Technologies, and as solid support for RNA synthesis, polystyrene Universal Support III from Eurogentec was used. Oligonucleotide synthesis was performed on a 1000 nmol scale. 8-Br-dG-phosphoramidite (BrG) was purchased from Eurogentec, and all other phosphoramidites were purchased from Hongene Biotech Germany GmbH. 2’-OMe-8-Br-G phosphoramidite (BrGm) was synthesised as described in the Electronic Supplementary Information. All phosphoramidites were used as cyanoethyl-diisopropyl-phosphoramidites (with a 2’-TBS protection group for RNA phosphoramidites). The following protection groups for nucleobases were used: dC(Ac), dG(iBu), 8-Br-dG(dmf), rG(iBu), rC(Ac). All phosphoramidites were used in a concentration of 0.1 M in acetonitrile.

The following reagent mixes were used during solid support synthesis: acetonitrile (99.9%, extra dry over molecular sieves from Thermo Scientific); tetrahydrofuran/2,6-lutidine/acetic anhydride 8/1/1 (v/v/v) (Cap A); tetrahydrofuran/N-methylimidazole 84/16 (v/v) (Cap B); tetrahydrofuran/water/pyridine/iodine 77/2/21/2,54 (v/v/v/w) (Oxidiser); dichloromethane/trichloroacetic acid 97/3 (v/w) (Deblocking Mix). HPLC grade tetrahydrofuran, HPLC grade dichloromethane, and pyridine were purchased from Fisher Scientific, N-methylimidazole was purchased from Carbolution, 2,6-lutidine was purchased from TCI, acetic anhydride was purchased from Carl Roth, and trichloroacetic acid was purchased from Sigma Aldrich. Pre-mixed 0.25 M 5- (ethylthio)-1H-tetrazole in acetonitrile from Sigma-Aldrich was used as an activator.

The following reaction sequences were used for oligonucleotide synthesis: detritylation, coupling (2 min for DNA, 10 min for RNA and all unnatural building blocks), capping, oxidation, and capping. Capping was performed twice to guarantee complete capping of free OH-groups after coupling. The synthesis was carried out DMT-on.

After synthesis, the DNA oligonucleotides were cleaved from CPG solid support and deprotected by adding 1 mL of 28% NH3 and shaking at 55 °C for 16 h. After cooling to room temperature, the solution was diluted with 1 mL of aqueous NaCl solution (100 mg/mL) and purified with Glen-Pak™ DNA purification cartridges from Eurogentec according to the manufacturer’s standard protocol. The RNA oligonucleotides were cleaved from the polystyrene Universal Support III by adding 0.5 mL of 2 M NH3 in MeOH and shaking at 25 °C for 1 h. Then, the NH3 in MeOH was partially evaporated by shaking at 35 °C for 1.5 h before adding 1 mL of 28% NH3 to the residue and shaking at 55 °C for 16 h. After that, NH3 was evaporated again by shaking at 50 °C for 5 h and finally freeze-dried overnight. The residue was redissolved in 115 µL of DMSO, and 2’-TBS deprotection was carried out by adding 60 µL of triethylamine and 75 µL of triethylamine trihydrofluoride complex and shaking at 65 °C for 2.5 h. After cooling to room temperature, the solution was diluted with Quenching Buffer from Eurogentec, and purification was performed with Glen-Pak™ DNA purification cartridges from Eurogentec according to the manufacturer’s standard protocol.

Concentrations of the elution fractions were calculated from the A260 values measured with a Denovix DS-11 spectrophotometer, and the sequence absorbance was determined at https://oligocalc.eu/. Extinction coefficients of modified guanosines were assumed equal to those of natural guanosines. The obtained oligonucleotides were analysed via LC-ESI-MS (see Electronic Supplementary Information).

### 2.3 Brominated RNA building block and synthesis of Z-RNA

The left-handed Z-helix is thermodynamically unfavoured due to increased steric strain and electrostatic repulsions in the phosphate backbone. Stabilising the Z-helix with its alternating *anti*-C/*syn*-G sequence requires chemically modified guanosine building blocks that favour the *syn*-conformation. This can be achieved by *C*8-substitution, e.g., with bromine (Br) or methyl (Me). (Krall et al., 2023)

While the phosphoramidite building block of 8-Br-dG (BrG) for Z-DNA was obtained commercially, the building block of 2’-OMe-8-Br-G phosphoramidite (BrGm) for Z-RNA stabilisation was synthesised in three steps starting from N-isobutyryl-2’-O-methylguanosine (1) according to Figure 1. The 2’-methoxy substitution was chosen because it allows a shorter synthetic route compared to a natural 2’-hydroxy group, which must be protected as a silyl ether. The ^Br^G _m_ phosphoramidite was used to synthesise the stabilised Z-RNA 5’-rC ^Br^G_m_ rC ^Br^G_m_ rC rG-3’. Detailed synthetic procedures and analytical data are given in the Electronic Supplementary Information.

**Figure 1.**
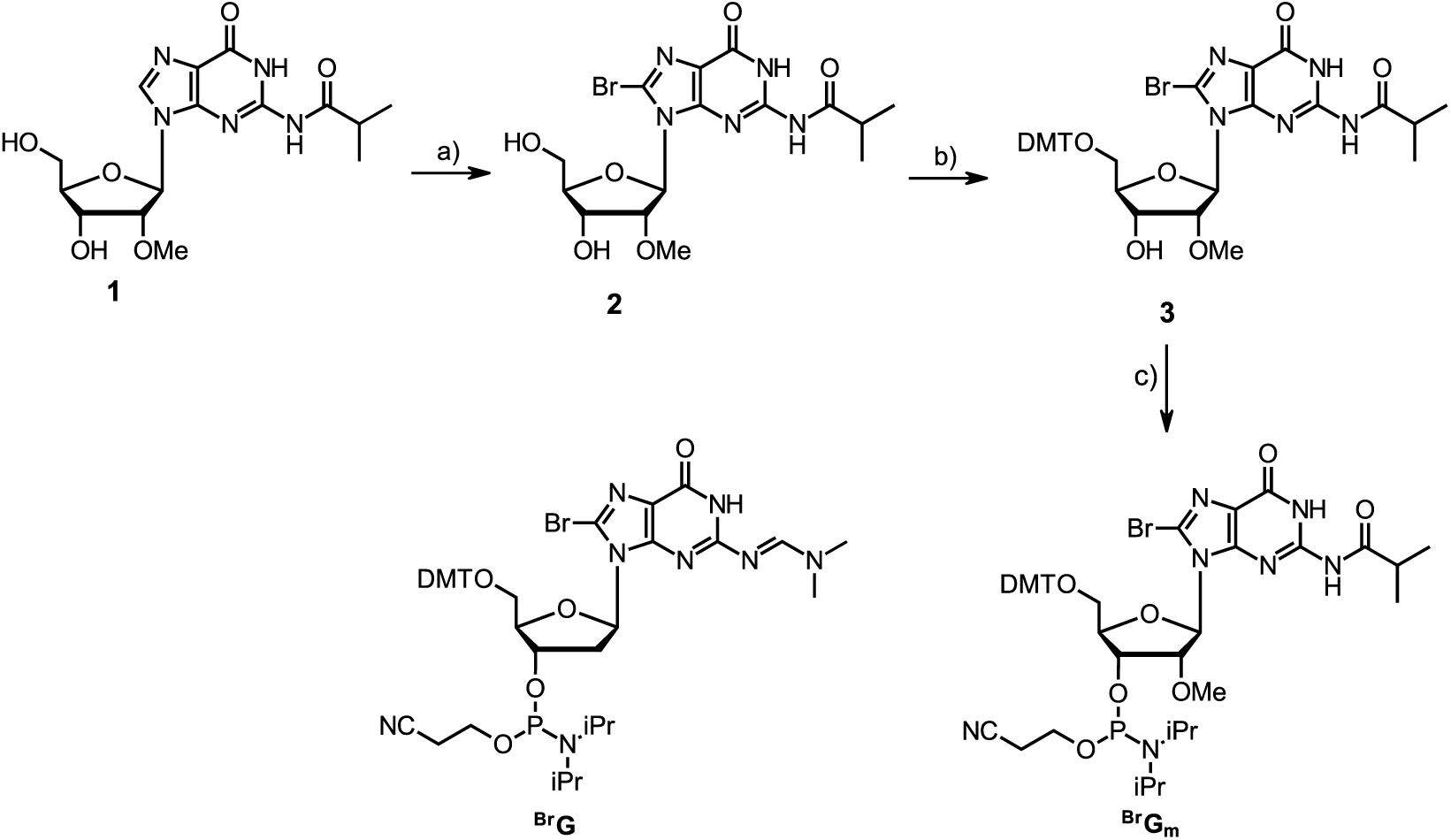
Structures of 8-Br-dG phosphoramidite (^Br^G) and 2’-OMe-8-Br-G phosphoramidite (^Br^G_m_) with synthesis; a) NBS, MeCN/H_2_O 2:1, r.t., 23 h, 90%; b) DMTCl, NEt_3_, DMAP, pyridine, r.t., 2.5 h, 94%; c) 2-cyanoethylphosphor-*N,N,N’,N’*-tetraisopropylbisamidite, tetrazole, CH_2_Cl_2_, 0 °C – r.t., 3.5 h, 89%.

### 2.4 Preparation of synthetic G4-NA and Z-NA

DNA and RNA oligos (Table 1) were annealed at 90 °C (3 min) and gradually cooled down to 30°C (over 2 h) in Z-buffer (25 mM Tris, pH 6.0, with 6.25 mM CaCl_2_ and 1 mM MgSO_4_ for Z-NA stabilisation) and G4-buffer (10 mM Tris, pH 7.5, with 100 mM KCl for G4-NA stabilisation). The concentration of oligos was 1.92 µM for DNA staining and 5 µM for circular dichroism measurements.

### 2.5 Fluorescence measurements of synthetic oligos using DNA-binding dyes

The annealed oligos (48 µL, 2 µM) were mixed with DNA-binding dyes (2 µL, 50 µM) in a 384-well plate (Corning, black), resulting in a working concentration of 1.92 µM oligos and 2 µM dye. For SYTOX™ Red binding and detection, 35 µL of the 2 µM oligos were mixed with 15 µL of 5 µM SYTOX™ Red, resulting in 1.4 µM oligos and 1.5 µM dye. Fluorescence spectra were recorded in a PerkinElmer EnSight multimode plate reader using the following settings (Table 2). The average fluorescence values (recorded over a 100 nm range emission spectra around the fluorescence peak) were then quantified for every dye after subtracting the value of the blank dye sample. The excitation wavelength must be at least 20 nm lower than the emission spectrum to avoid collection of excitation light by the detector. We therefore used excitation wavelengths slightly below the excitation maximum for each dye, resulting in lower total fluorescence in exchange for collecting the entire emission spectrum for each dye.

**Table 2.** PerkinElmer EnSight multimode plate reader settings for fluorescence quantification of dye/NA complexes.

| *Dye* | *Excitation* | *Emission spectrum* | *Gain* |
| --- | --- | --- | --- |
| **TOTO™-1** | 470 nm | 510-610 nm | 1000 |
| **TOTO™-3** | 610 nm | 630-730 nm | 1000 |
| **YOYO™-1** | 470 nm | 490-590 nm | 100 |
| **YOYO™-3** | 580 nm | 600-700 nm | 1000 |
| **BOBO™-1** | 440 nm | 460-560 nm | 100 |
| **BOBO™-3** | 550 nm | 570-670 nm | 100 |
| **POPO™-1** | 400 nm | 430-530 nm | 50 |
| **POPO™-3** | 510 nm | 530-630 nm | 100 |
| **SYTOX™ Green** | 470 nm | 500-600 nm | 100 |
| **SYTOX™ Red** | 580 nm | 600-700 nm | 1000 |
| **PI** | 550 nm | 570-670 nm | 1000 |

### 2.6 Circular Dichroism

The structure of NA was confirmed by far-UV circular dichroism (CD) spectra measured on a JASCO J-810 spectropolarimeter (JASCO Corp., Tokyo, Japan) within the wavelength range of 220–320 nm, data pitch at 0.5 nm, scanning speed at 100 nm/min, response at 4 sec, and bandwidth at 2 nm. The measurements were performed using 50 μL of the sample in a cuvette with a 0.3 cm optical path. Samples of 5 μM DNA and RNA were prepared in 1 × Z-buffer or 1 × G4-buffer. Signal (ellipticity) was measured in millidegrees.

### 2.7 Biofilm preparation

The bacterial strain used in this study was the 29213 *Staphylococcus aureus* (*S. aureus*) reporter strain with a plasmid (pKK30) containing a resistance gene for trimethoprim (TMP) stored at -80 °C in 15 % glycerol. Single colonies were grown on tryptic soy agar with 10 µg/mL TMP and stored at 4-8 ^°^C for up to 2 weeks. A single colony was inoculated into 20 mL tryptic soy broth (TSB) (MP Biomedicals*, #157152, Lot # U1123052160-1*) with 10 µg/mL TMP and 0.2 M NaCl (Sigma-Aldrich, #S5886) in a 100 mL sterile Erlenmeyer flask. The overnight cultures were incubated at 37 °C, 180 RPM.

The bottom of a 96-well plate (*Ibidi, #89626, IbiTreat*) was coated with 10 % human plasma in TSB for 30 min, aspirated, and replaced with 200 µL *S. aureus* overnight culture diluted 1:1000 in TSB + 0.2 M NaCl + 10 µg/mL TMP. The plate was incubated statically at 37 °C for 2 h to allow bacterial adhesion before replacing the medium and incubating for 2 days at 37 °C, 180 RPM.

### 2.8 Visualisation of eNA by immunofluorescence labelling

Primary mouse anti-B-DNA antibody AB1 (*Abcam*) or mouse anti-dsRNA antibody J2 (*Jena Bioscience*) was labelled using the antibody labelling kit from ProteinTech (*Flexable, #KFA546*) for labelling IgG2a mouse antibody with AlexaFluor 405 following the manufacturer’s instructions.

The 2-day biofilms were washed twice with PBS (*VWR, #E703*), then blocked with 3 % BSA (*Sigma-Aldrich, # A7906*) in PBS for 15 min before adding a mix of Z22 (diluted 1:100), BG4 (diluted 1:100), and AB1 antibody (diluted 1:100), or anti-dsRNA antibody (diluted 1:100) in 3% BSA. After 2 h incubation at RT, the unbound antibody was washed off with 3 % BSA twice. The membrane-binding dye FM 4-64 (10 mg/L) was added to some samples to visualise bacteria. Biofilms were visualised with a Zeiss LSM 900 Airyscan2 confocal laser scanning microscope (CLSM) (*Carl Zeiss, Germany*) using a 63x 1.4 NA Plan-Neofluoar objective and the Airyscan imaging mode. Excitation and emission wavelengths are listed in Table 3. Images were prepared in Zen.

**Table 3.** CLSM settings used for visualising eNA in various Figures.

| Figure, Panel | Dye | Excitation (nm) | Emission (nm) |
| --- | --- | --- | --- |
| Figure 3, B | FM 4-64 | 488 | 615-700 |
| Figure 3, C | BG4 | 639 | 620-700 |
| Figure 3, D | Z22 | 488 | 450-560 |
| Figure 3, E | AB1 | 405 | 400-505 |
| Figure 3, G | FM 4-64 | 488 | 495-700 |
| Figure 3, H | J2 | 405 | 400-520 |
| Figure 4, A+B | YOYO™-3 | 555 | 565-700 |
| Figure 4, A | SYTO™ 41 | 405 | 400-575 |
| Figure 4, C | BOBO™-3 | 555 | 450-700 |
| Figure 4, D | POPO™-3 | 555 | 450-700 |
| Figure 4, E | TOTO™-3 | 639 | 450-700 |
| Figure 4, F | TOTO™-1 | 488 | 450-700 |
| Figure 4, G | SYTOX™ Red | 639 | 620-700 |
| Figure 4, H | SYTOX™ Green | 488 | 490-575 |
| Figure 4, I | PI | 555 | 450-700 |
| Figure 4, J | TSA | 488 | 490-700 |
| Figure 5, E | TOTO™-3 | 639 | 450-700 |
| Figure 5, F | TOTO™-1 | 488 | 450-620 |
| Figure 6, A+B | YOYO™-3 | 555 | 580-700 |
| Figure 6, C | TOTO™-1 | 488 | 450-595 |
| Figure 6, D+E | BOBO™-3 | 555 | 590-700 |
| Figure 6, F | TOTO™-1 | 488 | 450-580 |
| Figure 6, G+H | POPO™-3 | 555 | 570-700 |
| Figure 6, I | TOTO™-1 | 488 | 450-555 |
| Figure 6, J+K | PI | 555 | 585-700 |
| Figure 6, L | TOTO™-1 | 488 | 450-575 |
| Figure 7, A-F | Labelled ATP | 639 | 400-700 |
| Figure 7, A-C | SYTO™ 41 | 405 | 400-700 |

### 2.9 Visualisation of eNA by DNA-binding dyes

The 2-day biofilms were washed twice with PBS before adding dyes. 5 µM SYTO™ 41 was added as an intracellular dye, 10 mg/L FM 4-64 was added as a membrane dye, and 2 µM SYTOX™ Red, SYTOX™ Green, PI, TOTO™-1, TOTO™-3, BOBO™-3, POPO™-3, or YOYO™-3 diluted in PBS was added to the wells individually or in a combination of eNA dyes. After 20 min of incubation at room temperature, the wells were inspected under the microscope with settings listed in Table 3.

### 2.10 Visualisation of extracellular G4s by tyramide signalling amplification

5 µM hemin (Sigma-Aldrich, #H9039) (from a fresh 2 mM stock, dissolved in 10 % NaOH in milliQ water, kept away from light) was added to overnight cultures, and biofilms were otherwise prepared as described above, with media supplemented with 5 µM hemin. 2-day biofilms were washed twice with modified MES buffer (25 mM MES (Sigma-Aldrich, #M3671), 0.2 M NaCl, and 0.01 M KCl (Sigma-Aldrich, #1.04936)) adjusted to pH 6.5. The MES buffer was gently removed, and 90 µL tyramide reagent (1:100 dilution, Alexa Fluor 488-conjugated Tyramide (Invitrogen, #B40953), 2 mM ATP (ThermoFischer, #R0441), and 0.005 % hydrogen peroxide (Sigma-Aldrich, #216763) in modified MES buffer was added and incubated for 90 minutes (37 °C, 50 RPM). After incubation, the tyramide reagent was aspirated, and the biofilms were washed twice in MES buffer. The biofilms were subsequently blocked with 3 % BSA in PBS for 15 minutes at room temperature before adding BG4 (diluted 1:100 in 3 % BSA). After 2 h at room temperature, the biofilms were washed twice with 3 % BSA to remove unbound mAbs. The biofilms were visualised under a CLSM microscope (settings listed in Table 3).

### 2.11 PolyA-tailing for visualisation of eRNA

The 2-day biofilms were washed twice with PBS. Phosphatase was added to remove potential phosphates at the 3’-OH end of RNA and incubated at 37 °C for 10 min. Phosphatase-treated biofilms were washed twice with PBS. A PolyA-tailing kit (ThermoFischer, #AM1350) mix, in which 4 x lower ATP levels than recommended by the manufacturer and 1/3 of the ATP was replaced with ATP labelled with ATTO-647N (Jena Bioscience, #NU-807-647N) (Weissinger et al., 2025), was added to biofilms and incubated for 20 min at 37 °C. Unbound ATP was washed off by washing with PBS twice. The bacteria were stained intracellularly with 5 µM SYTO™41.

## 3. RESULTS

### 3.1 Turn-on fluorescence when interacting with annealed DNA and RNA oligos

We first assessed the fluorescence intensity of the dyes when bound to DNA or RNA *in vitro*, using annealed synthetic oligos folded into secondary structures B-DNA, Z-DNA, G4-DNA, A-RNA, Z-RNA, and G4-RNA (Figure 2). While B-DNA, A-RNA, G4-DNA, and G4-RNA sequences for staining experiments could be purchased, Z-DNA and Z-RNA were synthesised. The structures of B-DNA, A-RNA, Z-NA, and G4-NA were confirmed using circular dichroism (CD) (Figure S1). While monobromated DNA revealed a strong transition from B- to Z-DNA, it was not sufficient for RNA, which still resembled an A-RNA shape. Furthermore, dibrominated DNA showed an additional change in CD, while dibrominated RNA’s CD had its most indicative change associated with the transition from A- to Z-RNA. We, therefore, have chosen dibrominated DNA as well as RNA as substrates for annealing Z-NA.

**Figure 2.**
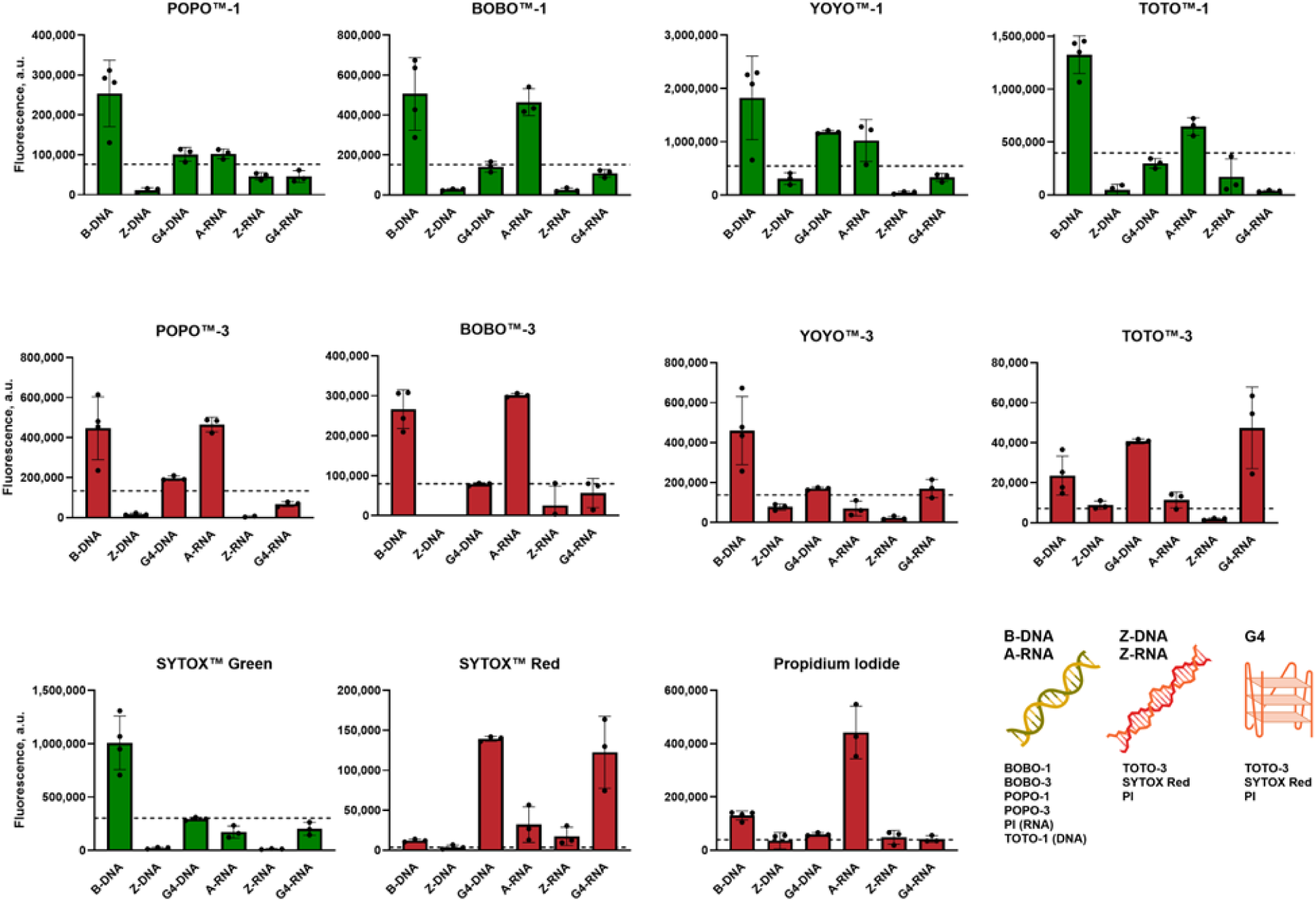
Fluorescence intensity after interaction with annealed DNA and RNA oligos. In each panel, all oligos are measured with the same settings. Bar graphs show mean values (n=3), error bars = S.D. The dotted line indicates a value of 30% fluorescence compared with the signal from B-DNA. We proposed this value as a cut-off in the assessment of whether other eNA structures could be detected by the same dye. Note that the concentration of dyes was 2 µM and the concentration of synthetic eNA structures was 1.92 µM in all mixtures, except for SYTOX red (1.4 µM dye and 1.5 µM eNA structures). Cartoons of eNA structures list the dyes that can detect them. See Table 2 for the spectra acquisition settings, Table S1 for the absolute fluorescence values, and Figure S2 for the full spectra.

Each dye was assessed separately (see full spectra in Figure S2), and the average fluorescence intensities (see Table 2 for the wavelength ranges used for averaging the intensity signals) for a specific dye were compared for the six different NA structures (Figure 2, Table S1). All dyes increased in fluorescence when bound to B-DNA, but their fluorescence intensity when bound to non-canonical nucleic acids compared to B-DNA was highly variable. There was a distinct pattern in the properties of green- vs. red-fluorescent dyes. The green-fluorescent cyanine dimers (except BOBO™-1) showed the strongest fluorescence when bound to B-DNA. In contrast, red-fluorescent cyanine dimers POPO™-3 and BOBO™-3 showed strong fluorescence when bound to either B-DNA or A-RNA. Other dyes showed an even higher fluorescence when bound to non-canonical structures compared to B-DNA. These were TOTO™-3 and SYTOX™ Red, which showed the strongest fluorescence when interacting with G4-NA. Moreover, PI displayed a 3-fold higher fluorescence when binding to A-RNA compared to B-DNA. Little or no fluorescence was observed from all dyes when interacting with Z-NA.

Dye molecules can stack and form aggregates emitting red-shifted fluorescence (Ichimura, 1975). TOTO™-1 and POPO™-1 produced red-shifted fluorescence in Z-RNA, unlike Z-DNA (Figure S2). A similar phenomenon was reported for single-stranded RNA, where the acridine orange dye bound anionic phosphate groups, resulting in dye stacking and high red-shifted fluorescence (Ichimura, 1975).

Overall, these data show specificity for green-fluorescent POPO™-1, BOBO™-1, YOYO™-1, TOTO™-1, and SYTOX™ Green toward B-DNA and A-RNA, while G4 can be detected by red-fluorescent TOTO™-3, YOYO™- 3, and SYTOX™ Red, and A-RNA can be detected by BOBO™-1, BOBO™-3, POPO™-3, and PI.

### 3.2 CLSM images of biofilms confirm the detection of non-canonical eNA by the far-red dyes

We next proceeded to assess the dyes for microscopy imaging. In Figure 3, we validate our biofilm model by immunolabelling, showing the presence of G4-NA (Figure 3C), Z-NA (Figure 3D), B-DNA (Figure 3E), and A-RNA (Figure 3H) associated with the surface of *S. aureus* cells.

**Figure 3.**
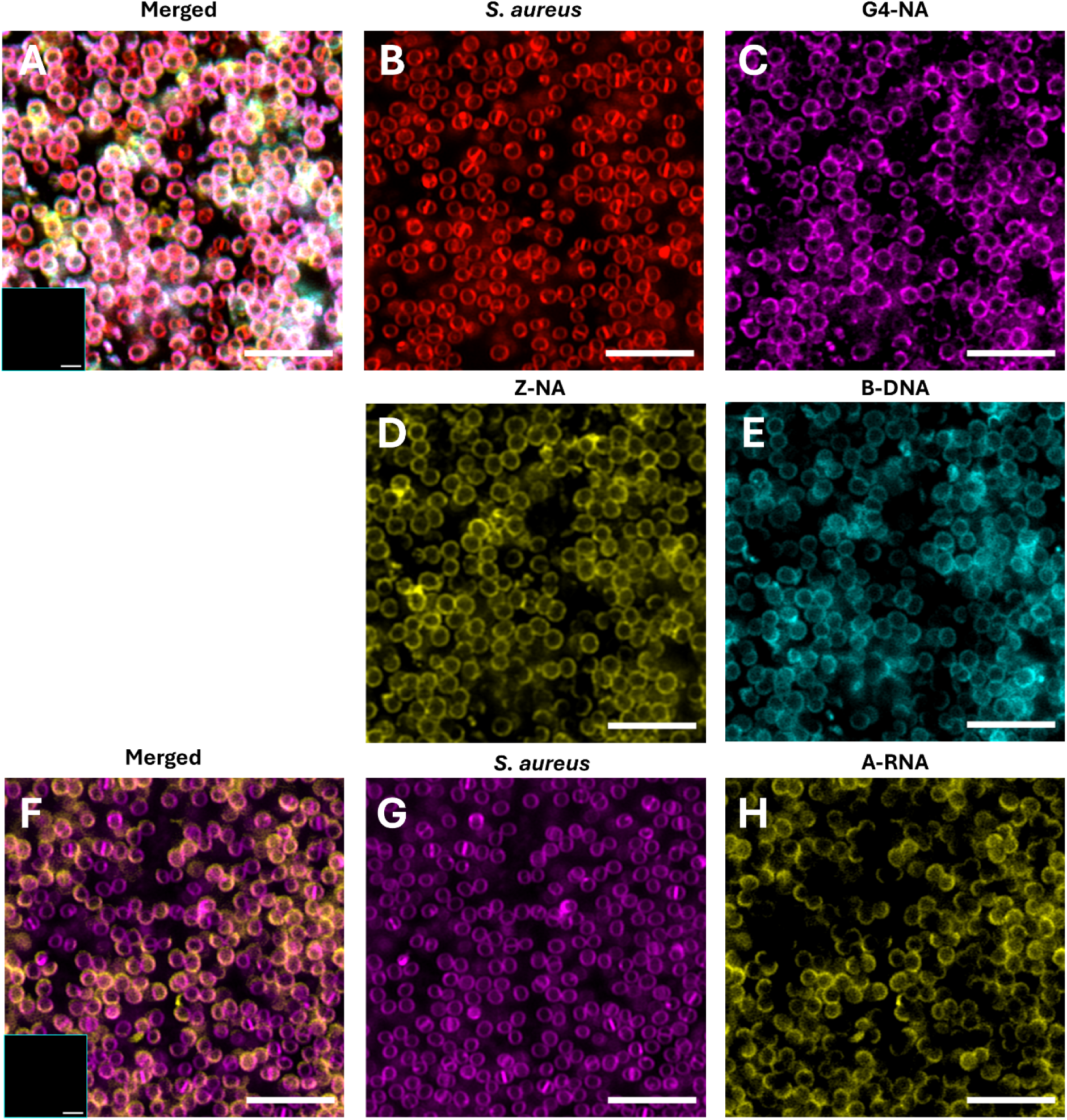
2-day *S. aureus* biofilms hold abundant amounts of plasma membrane-associated eNA. CLSM images of immunolabelled 2-day biofilms. Sample 1 (A-E): **A** merged + unstained control in the lower left corner, **B** *S. aureus* visualised with FM 4-64 (Red), **C** G-quadruplexes visualised with BG4 (Magenta), **D** Z-NA visualised with Z22 (Yellow), and **E** B-DNA visualised with AB1 (Cyan). Sample 2 (F-H): **F** merged + unstained control in lower left corner, **G** *S. aureus* visualised with FM 4-64 (magenta), **H** A-RNA visualised with J2 (Yellow). Scale bars = 5 µm.

After confirming the presence of non-canonical eNA structures, we evaluated the dyes of interest. First, we confirmed the presence of cells with the membrane-permeant SYTO™41, shown in combination with the extracellular YOYO™-3 in Figure 4A. Next, we evaluated each dye alone. YOYO™-3 (Figure 4B), BOBO™-3 (Figure 4C), POPO™-3 (Figure 4D), and TOTO™-3 (Figure 4E) resulted in surface-associated fluorescence, similar to observations by immunofluorescence labelling. PI, which had shown a strong preference for A-RNA oligos (Figure 2), only stained intracellular NA of dead cells (Figure 4I). However, some extracellular staining was visible when increasing the histogram drastically (Figure S3C), consistent with immunolabelling results using the A-RNA-specific antibody J2 (Figure 3H). SYTOX™ Red stained all cells intracellularly (Figure 4G), indicating that it permeabilised the membrane. SYTOX™ Red was supplied in DMSO at a low concentration, which made the DMSO concentration lethal to the bacteria. Therefore, it is not suitable for microscopy of biofilms. SYTOX™ Green and TOTO™-1, for which our initial analyses had shown a strong preference for B-DNA (Figure 2), did not stain eNA in the sample even when increasing the histograms drastically (Figure S3A+B). Both dyes only stained intracellular NA of dead cells (Figure 4H+F), indicating that B-DNA constitutes a minor fraction of the eNA.

**Figure 4.**
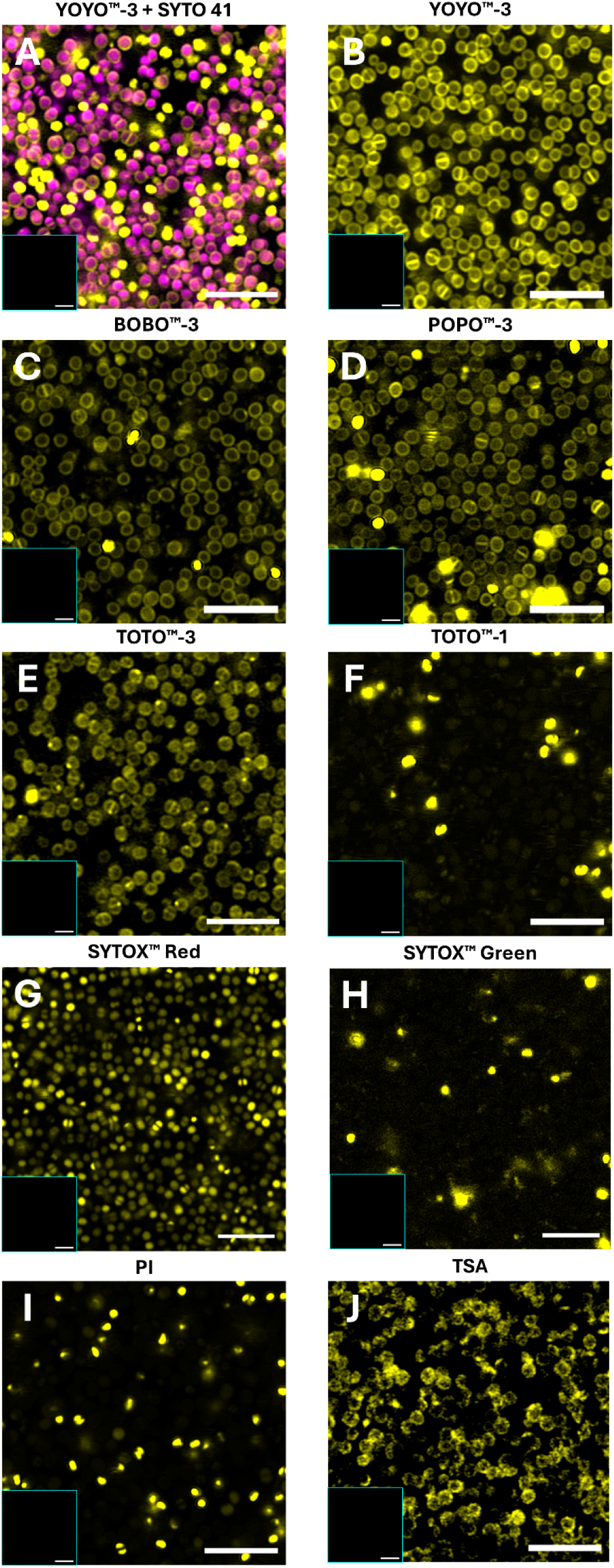
YOYO™-3, TOTO™-3, BOBO™-3, POPO™-3, and PI show potential for staining eNA in 2-day *S. aureus* biofilms. CLSM images of 2-day *S. aureus* biofilms visualised with **A** SYTO™ 41 (Magenta) and eNA visualised with **A** YOYO™-3 (Yellow), **B** YOYO™-3 (Yellow), **C** BOBO™-3 (Yellow), **D** POPO™-3 (Yellow), **E** TOTO™-3 (Yellow), **F** TOTO™-1 (Yellow), **G** SYTOX™ Red (Yellow), **H** SYTOX™ Green (Yellow), **I** PI (Yellow), **J** TSA (Yellow). The unstained control in lower left corner of all images. Scale bars = 5 µm.

We also used tyramide signal amplification (TSA) as an alternative method to visualise G4 via its peroxidase activity when complexed with hemin (Figure 4J). TSA showed extracellular signal associated with the surface of *S. aureus*, similar to the signal observed using G4-specific immunolabelling (Figure 3C). It should be noted that the catalytic activity of G4/hemin varies depending on the type of G4 structure and the sequence of neighbouring nucleotides. A negative TSA signal, therefore, does not exclude the presence of G4. If the aim is to visualise G4, we therefore conclude that TOTO™-3 or YOYO™-3 is most suited. It should be noted that several small-molecule dyes have been developed for G4-specific detection, including N-methyl mesoporphyrin IX (NMM) (Arthanari et al., 1998), thioflavin T (Bradford et al., 2024; Xu et al., 2016), N,N’-diethylthiacarbocyanine iodide (DTC) (Kerwin et al., 2001), TOR-G4 (Lumiprobe), and BioTracker QUMA-1 (Sigma-Aldrich). However, these dyes are all cell-permeable, which makes them unsuitable for this study.

### 3.3 Combination of green- and red-fluorescent dyes to detect non-canonical eNA

The different fluorescence properties of green and red fluorescent dyes might provide an opportunity to distinguish non-canonical structures from B-DNA in a complex sample by using the dyes in combination. To test this hypothesis, we combined TOTO™-1 and TOTO™-3. Emission spectra confirmed that TOTO™-1 detects B-DNA and A-RNA (Figure 5), but TOTO™-1 fluorescence was approximately halved when used in combination with TOTO™-3 (Table S1), indicating that the dyes compete for binding sites. In comparison to TOTO™- 1, the intensity of TOTO™-3 was relatively low when binding to B-DNA, A-RNA, or G4-DNA (Figure 5A, B). However, TOTO™-3 fluorescence was more prominent compared to TOTO™-1 when binding to G4-RNA or to Z-NA, although overall fluorescence intensity from Z-NA was low. These results highlight complex interactions of DNA-binding dyes with different nucleic acid structures and corroborate that G4-RNA and possibly Z-NA can be distinguished from B-DNA in a complex sample by combining TOTO™-1 and TOTO™-3. Indeed, CLSM imaging with the dye combination showed that TOTO™-1 primarily stained intracellular NA in dead cells, while TOTO™-3 stained eNA (Figure 5D-F), indicating that the G4-NA detected by immunolabelling might be G4-RNA.

**Figure 5.**
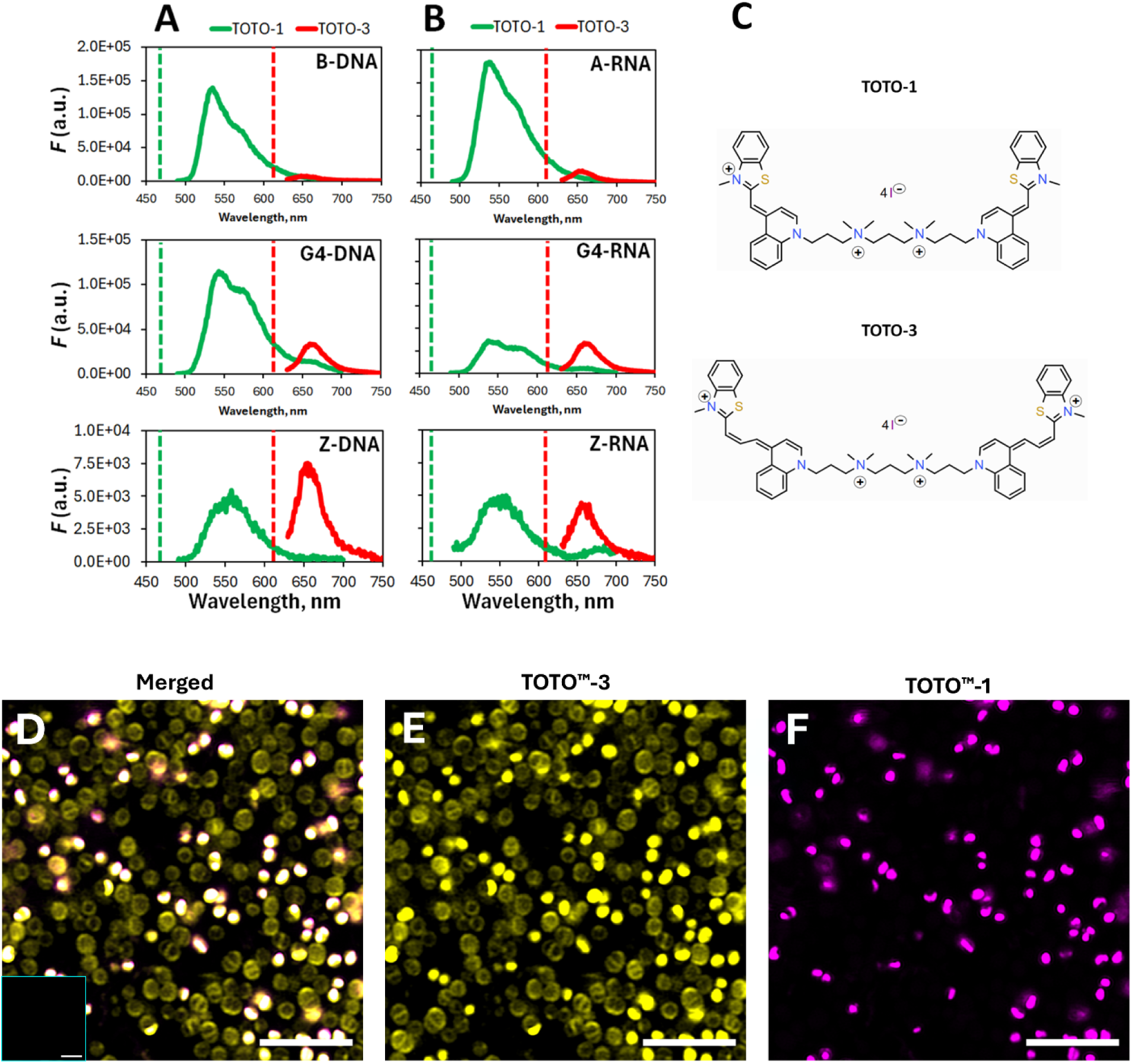
Combination of TOTO™-1 and TOTO™-3 with different specificity for B-DNA and non-canonical structures. **A+B** Fluorescence emission spectra recorded for DNA and RNA complexes with TOTO™-1 and TOTO™-3 in equimolar concentrations (n = 3 for each combination). Vertical dotted lines indicate excitation wavelengths at 470 nm (TOTO™-1) and 610 nm (TOTO™-3). **C** Chemical structures of TOTO™-1 and TOTO™-3. **D-F** CLSM images of 2-day *S. aureus* biofilms stained with TOTO™-3 (Yellow) and TOTO™-1 (Magenta) showing the merged image and the individual stains. The unstained control in the lower left corner. Scale bars = 5 µm.

We subsequently tested combinations of TOTO™-1 with either YOYO™-3, BOBO™-3, POPO™-3, or PI (Figure 6). Again, TOTO™-1 only stained intracellular NA in dead cells, whereas YOYO™-3, BOBO™-3, and POPO™- 3 also stained the surface-associated eNA we previously detected by antibodies (Figure 6). PI only showed a very dim signal from the surface-associated eNA, but a strong intracellular signal from dead cells. Surprisingly, these dead cells were invisible to TOTO™-1, indicating that PI competes with TOTO™-1 for binding (they are both intercalating dyes), or that PI quenches TOTO™-1 fluorescence.

**Figure 6.**
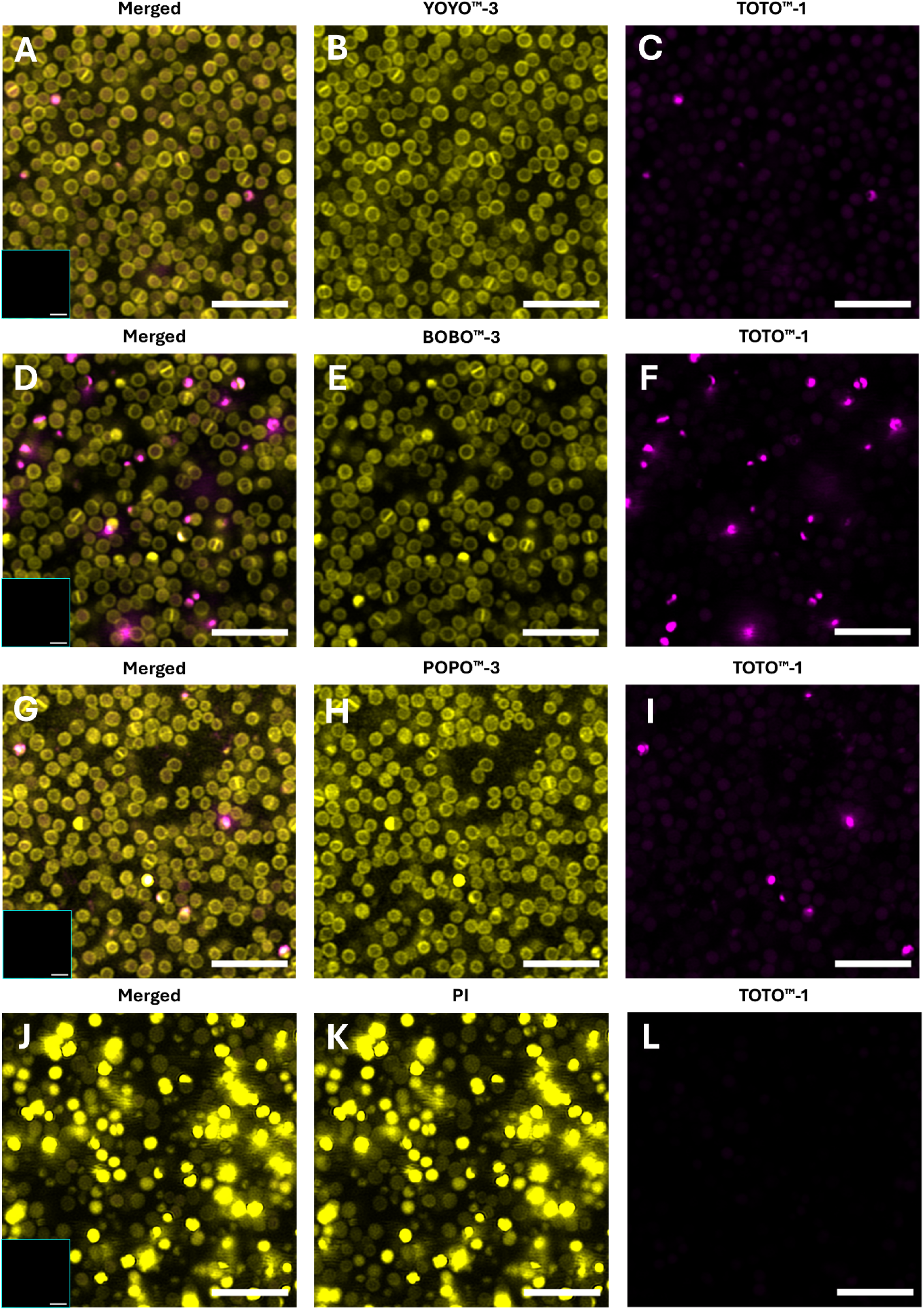
Combination of NA-binding dyes with different specificities for B-DNA and non-canonical structures. CLSM images of 2-day *S. aureus* biofilms. **A-C** TOTO™-1 **+** YOYO™-3 staining showing the merged image and individual dyes. **D-F** TOTO™-1 **+** BOBO™-3 staining showing the merged image and individual dyes. **G-I** TOTO™-1 **+** POPO™-3 staining showing the merged image and individual dyes. **J-L** TOTO™-1 **+** PI staining showing the merged image and individual dyes. The unstained control in the lower left corner of the merged images. Scale bars = 5 µm.

Lastly, we observed less eNA fluorescence if dyes were combined with the membrane stain FM 4-64 (data not shown). This prompted us to investigate whether FM 4-64 impacts the fluorescence signal from nucleic acidbinding dyes directly, using our prefolded oligos (Table 1). Indeed, FM 4-64 reduced the signal from e.g., YOYO™-1, particularly when bound to G4-DNA (Figure S4), most likely due to displacement of the dye, indicating that FM 4-64 binds to the hydrophobic G4 but does not light up when bound. FM 4-64 staining should therefore be avoided when detecting G4-NA in biofilms.

### 3.4 RNA visualisation

Our results indicate that the eNA associated with the plasma membrane of bacteria in our *S. aureus* biofilms is not only DNA but also RNA. The J2 antibody, with specificity for A-RNA, confirmed the association of A- RNA with the cell surface (Figure 3H), and it is known that the antibodies used to detect Z-NA and G4 have affinity for both the DNA and RNA versions of these structures, which is also indicated by the fact that Z-NA (Z22) and G4-NA (BG4) did not always co-localise with B-DNA (AB1) (Figure 3). We lack methods that strictly detect RNA besides the J2 antibody, which only detects A-RNA. To address this, we adapted a novel approach to use labelled ATP and polyA-tailing to enable imaging of RNA (Weissinger et al., 2025) in the biofilm matrix.

Without phosphatase pre-treatment, the fluorescence from polyA-tailing was mostly localised intracellularly, presumably in dead cells (Figure 7A). When the biofilm was treated with phosphatase to enable extracellular polyA-tailing by removing inhibitory 3’-terminal phosphorylation, fluorescence was detected around bacterial cells (Figure 7B+E, indicated with arrows), localising where we previously detected A-RNA with fluorescence immunolabelling (Figure 3H). The control without Poly(A) Polymerase I (*E*-PAP) showed no labelled ATP, ruling out unspecific incorporation of ATP in the biofilm. Thus, we demonstrate polyA-tailing as a novel method for RNA-detection technique in biofilms.

**Figure 7.**
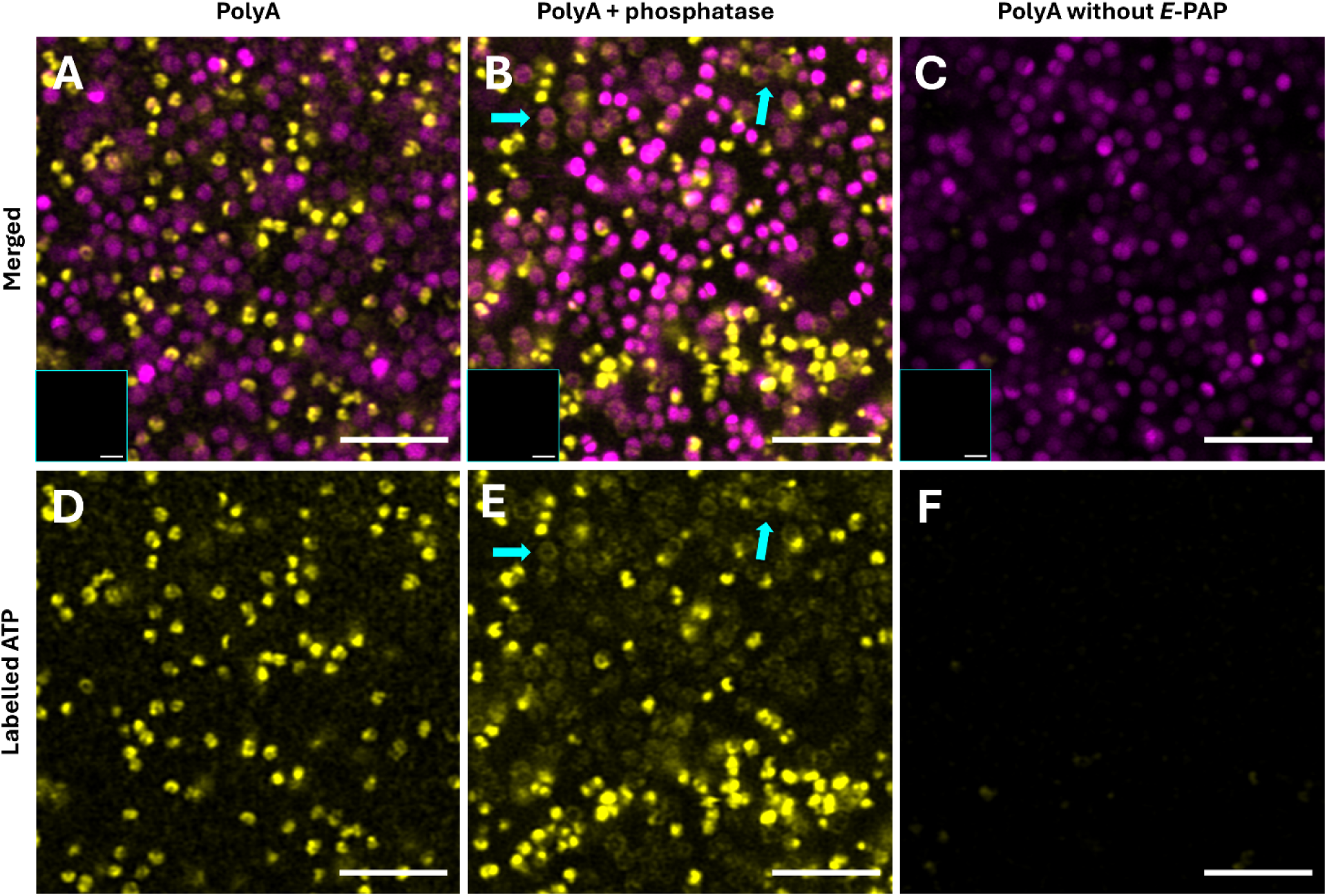
PolyA-tailing with labelled ATP shows detectable visualisation of RNA in 2-day biofilms when pre-treated with phosphatase. CLSM images of 2-day *S. aureus* biofilms. *S. aureus* visualised with SYTO™ 41 (Magenta) and labelled ATP (Yellow). **A + D** PolyA-tailing without a dephosphorylation step, **B + E** PolyA-tailing with a dephosphorylation step, **C + F** control without added *E*-PAP. Cyan arrows indicate ring-shaped staining patterns. **A-C** merged + unstained control in lower left corner. Scale bars = 5 µm.

## 4. DISCUSSION

Most biofilms contain eNA in the extracellular matrix, and the prevalence of non-canonical eNA structures is becoming increasingly evident. They provide emerging properties to the biofilm matrix, including extracellular electron transfer (Ajunwa & Meyer, 2026; Ajunwa et al., 2025; Kotoky et al., 2025; Saunders et al., 2020), peroxidase-like catalytic activity (Minero et al., 2024), and increased resistance to antibiotics (Chiang et al., 2013), the host immune system (Gallucci, 2024), and they can even be synthesised extracellularly, creating hubs of phase separation in biofilm (Minero et al., 2025). Therefore, visualising noncanonical eNA structures is critical for exploring their significance to biofilm biology more broadly.

This study aimed to determine if nucleic acid-binding dyes can detect non-canonical nucleic acid structures and possibly distinguish between B-DNA and these structures. We noticed a general pattern of green-fluorescent dyes being more selective for canonical structures (B-DNA and A-RNA). Among these dyes, TOTO™-1 and SYTOX™ Green showed the greatest preference for B-DNA, making them suitable for B-DNA-specific detection. In contrast, the red-fluorescent dyes exhibited broader specificity that included non-canonical structures. TOTO™-3 and SYTOX™ Red showed the most promising properties for the detection of non-canonical eNA. When bound to Z-DNA and Z-RNA, these were the only dyes that showed detectable fluorescence, albeit with lower intensity. Furthermore, TOTO™-3 and SYTOX™ Red detected G4-NA with up to 7-fold higher fluorescence intensity compared to B-DNA, making them promising candidates for G4 detection in biofilms.

Most surprisingly, the commonly used PI displayed 3-fold higher fluorescence intensity when bound to A-RNA compared to B-DNA. Although PI is widely used as an RNA-staining dye in other fields of research, e.g., in flow cytometry (Frankfurt, 1990), this finding has important implications for biofilm research, since PI is widely used for dead cell staining and eDNA visualisation. While eDNA is well described in biofilms, eRNA has received less attention (Mugunthan et al., 2025; Rosenberg et al., 2019). Our observations suggest that PI has significant potential for visualising eRNA in biofilm samples – possibly in combination with DNA-specific dyes.

In addition to the structural diversity, it should also be noted that eNA in biofilms interacts with a wide range of molecules that can impact staining results. For example, cationic polysaccharides such as chitosan can reduce the fluorescent intensity of dyes (Figure S5). Likewise, the DNA sequence and bacterial epigenetic changes, including differences in DNA methylation, may also influence the fluorescence of the dye (Minero et al., 2012). Staining of eNA in biofilms will thus be impacted by the variations in eNA sequence, structure, interactions, and the local environment.

### 4.1 Structural Basis for Differential Dye Binding to eNA

The dramatic difference in specificity between TOTO™-1 and TOTO™-3 provides insight into the structural requirements for non-canonical NA binding. The cyanine dimers TOTO™-1/-3 differ in the length of their unsaturated carbon chains (one versus three carbon atoms, respectively), making TOTO™-3 more hydrophobic and longer than TOTO™-1. This increased hydrophobicity appears critical for binding non-canonical structures. G4-NA is known to bind a plethora of hydrophobic molecules (Ma et al., 2020), and therefore, the hydrophobic properties of far-red dyes likely contribute to their increased binding affinity for G4-NA and consequent fluorescence enhancement, as was demonstrated for a far-red SYTO™60 (Lund, in press). Additionally, the longer hydrophobic linkers between the homodimers, e.g., in TOTO™-3, may make these dyes more geometrically compatible with both the elongated Z-NA structures and barrel-shaped G4-NA structures, both of which also tend to expose hydrophobic bases (Gray et al., 2014; Krall et al., 2023). In contrast, the less hydrophobic and shorter linker of TOTO™-1 appears optimised for intercalating into the regular B-DNA double helix, where base stacking provides a more uniform hydrophobic environment.

The molecular structure of SYTOX™ Red is not disclosed by the manufacturer (Thermo Fisher), making it difficult to explain mechanistically why this dye shows such versatile interaction with multiple NA conformations. Furthermore, its implementation for visualising eNA *in situ* is limited because it is supplied in low concentrations (5 µM) in DMSO, which makes it toxic to bacteria at the working concentration and results in intracellular staining (Figure 4I).

### 4.2 Validation and Application in Biofilms

To validate the presence and distribution of eNA structures in our biofilm model, we used antibodies specific for B-DNA (AB1), Z-NA (Z22), G4-NA (BG4), and A-RNA (J2). Immunofluorescence labelling revealed that all four eNA types were associated with the surface of *S. aureus* cells. These surface-associated eNA were detected by TOTO™-3, BOBO™-3, YOYO™-3, POPO™-3, PI, TSA labelling of G4/hemin peroxidase activity. In contrast, TOTO™-1 only stained the intracellular DNA of dead cells (Figure 5F), despite detection of surface-associated B-DNA by immunolabelling (Figure 3E). It is important to note that the *in vitro* and *in situ* experiments were conducted in different chemical environments, and differences in DNA sequence can also affect the binding and fluorescence intensity of the tested dyes. Such differences might therefore explain why TOTO™-1 was absent from the bacterial surface. However, the observation might also indicate that the eNA is predominantly RNA. The mAb Z22 and BG4 recognise both the DNA and RNA versions of their target structure, and AB1 has been reported by the manufacturer (Abcam) to show cross-reactivity with A-RNA.

Several evidence show that RNA is present on the surface of *S. aureus:* 1) absence of TOTO™-1 labelling in the presence of TOTO™-3 labelling (Figure 5), 2) positive immunolabelling with anti-A-RNA antibody J2 (Figure 3), and 3) positive signal from RNA-specific poly-A tailing (Figure 7D+E), which is highly specific for RNA. In the latter assay, RNases from lysed bacteria, including *S. aureus* (Numata et al., 2014), can leave 3’-phosphoryls (Bechhofer & Deutscher, 2019; Grünberg et al., 2021), which block the *E*-PAP. When we added a dephosphorylation step, we observed cell surface-associated RNA (Figure 7B + E), similar to the RNA detected by fluorescence immunolabelling (Figure 3H). The result confirms the presence of eRNA on the surface of *S. aureus* and indicates that RNases are released in these biofilms.

While this approach shows promise, the signals remained relatively weak compared to antibody labelling. Antibodies and intercalating dyes can bind and stain NA molecules many times, whereas the polyA-tailing technique solely marks the ends of RNA, which results in only staining an RNA molecule once, providing weak signals. Further optimisation of this technique could make it valuable for RNA detection in biofilm research.

## 5. CONCLUSION

The structural diversity of eNA in biofilms is not accounted for when imaging eNA in biofilms with conventional DNA-binding dyes, and non-canonical structures that can be abundant in biofilms are simply invisible to many dyes. We here fill an important methodological gap by re-evaluating DNA-binding dyes and their potential for binding other NA-structures than B-DNA.

TOTO™-3, YOYO™-1, or YOYO™-3 are suitable for broad eNA screening, detecting canonical and non-canonical structures simultaneously. We also show that the contrasting specificities of green- and red-fluorescent dyes can be exploited to distinguish between canonical and non-canonical structures in complex samples, e.g. using a combination of TOTO™-1 and TOTO™-3, respectively. However, one must expect lower overall fluorescence from each dye as they compete for binding sites, and results must subsequently be validated with more specific labelling techniques, such as fluorescence immunolabelling.

## 6. CRediT authorship contribution statement

**Freja Winther Sillesen** Conceptualisation, Methodology, Formal analysis, Investigation, Writing - Original Draft, Visualisation **Finn Dicke** Investigation, Formal analysis, Methodology, Writing – review & editing **Stephanie Kath-Schorr** Investigation, Formal analysis, Methodology, Writing – review & editing **Hannah Weissinger** Conceptualisation, Methodology, Investigation, Writing – review & editing **Jørgen Kjems** Conceptualisation, Resources, Writing – review & editing **Amalia Villum Jakobsen** Formal analysis, Investigation, Writing – review & editing **Daniel Otzen** Resources, Methodology, Writing – review & editing **Gabriel Antonio Salvador Minero** Conceptualisation, Methodology, Formal analysis, Investigation, Writing - Original Draft, **Rikke Meyer** Conceptualisation, Resources, Supervision, Project administration, Funding acquisition, Writing – review & editing.

## 8. ACKNOWLEDGEMENTS

We thank Dr. Brigitte Stadler for providing access to the plate reader. FWS, GASM, and RLM acknowledge the Villum Foundation (Grant no. 69632) and The Carlsberg Foundation (Grant no. CF23-1540) for funding a major part of this work. S.KS and F.D. thank the German Research Foundation (DFG), RTG 2550 and the UoC Forum for funding.

NMR spectroscopy was performed in the NMR Facility of the Department of Chemistry and Biochemistry at University of Cologne which benefits from generous funding by the German Research Foundation (Deutsche Forschungsgemeinschaft, DFG). The contribution of the NMR Facility is kindly acknowledged. Mass spectrometric characterization and data analysis of the compounds was performed in the MS facility of the Department of Chemistry and Biochemistry at the University of Cologne which benefits from generous funding by the German Science Foundation DFG. The contribution of the MS facility is kindly acknowledged.

