## Supplementary Information for "Visualising diverse extracellular nucleic acids in biofilms with DNA-binding dyes"

**Electronic Supplementary Information**

***Synthesis of ^Br^G_m_ phosphoramidite***

*N*-Isobutyryl-2’-*O*-methyl-8-bromoguanosine (**2**)

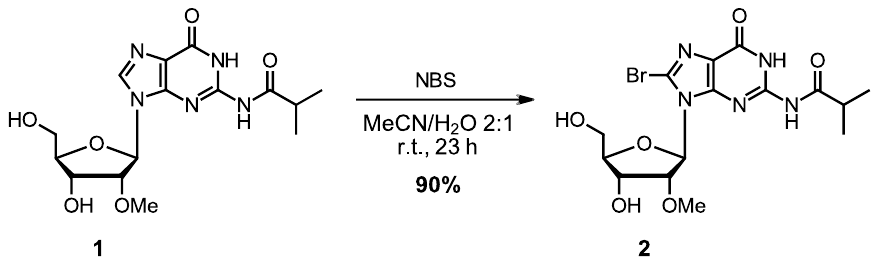

According to *Campitiello* (Campitiello et al., 2021) (1), *N*-isobutyryl-2’-*O*-methylguanosine (**1**, 1.50 g, 4.08 mmol, 1.00 equiv) was dissolved in 30 mL of H_2_O and 60 mL of MeCN. NBS (1.47 g, 8.28 mmol, 2.03 equiv) was added in three portions over 20 min and the reaction was stirred for 23 h until TLC showed full conversion of the starting material. The solvents were evaporated under reduced pressure, the residue was dissolved in MeOH, dry-loaded on celite and purified via flash column chromatography on SiO_2_ (CH_2_Cl_2_/MeOH 20:1 🡪 10:1). The product **2** was obtained as a colourless solid in a yield of 90% (1.65 g, 1.40 mmol).

| M(C_15_H_20_BrN_5_O_6_): | 446.26 g/mol. | | 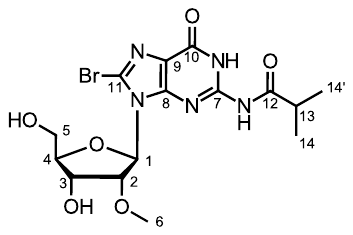 | |
| --- | --- | --- | --- | --- |
| Habitus: | Colourless solid. | |  |  |
| R_f_: | 0.40 (CH_2_Cl_2_/MeOH 20:1). | |  |  |
| ^1^H NMR: | (500 MHz, DMSO-*d*_6_) δ 12.20 (s, 1H, NH), 11.55 (s, 1H, NH), 5.87 (d, J = 6.1 Hz, 1H, H-1), 5.26 (d, J = 6.0 Hz, 1H, 3-OH), 4.87 – 4.79 (m, 2H, H-2, 5-OH), 4.39 (q, J = 5.3 Hz, 1H, H-3), 3.88 (q, J = 5.8 Hz, 1H, H-4), 3.67 (dt, J = 11.6, 5.8 Hz, 1H, H-5), 3.53 (dt, J = 11.6, 5.8 Hz, 1H, H-5'), 3.34 (s, 3H, H-6), 2.79 (hept, J = 6.8 Hz, 1H, H-13), 1.13 (d, J = 6.7 Hz, 6H, H-14, H-14'). | | | |
| ^13^C NMR: | (126 MHz, DMSO-*d*_6_) δ 180.3 (C-12), 153.6 (C-10), 149.9 (C-8), 148.1 (C-7), 123.8 (C-11), 121.0 (C-9), 87.6 (C-1), 86.0 (C-4), 79.2 (C-2), 68.7 (C-3), 61.6 (C-5), 57.7 (C-6), 34.8 (C-13), 18.89 (C-14 or C-14'), 18.87 (C-14 or C-14'). | | | |
| HR-MS (ESI): | Calc. | Exp. | | Dev. [ppm] |
|  | 446.06697 [M+H]^+^ | 446.06769 [M+H]^+^ | | +1.60 |
|  | 468.04892 [M+Na]^+^ | 468.04954 [M+Na]^+^ | | +1.34 |
| FT-IR (ATR): | 3443 (w), 3144 (w), 2932 (w), 2367 (w), 1697 (s), 1676 (s), 1603 (s), 1560 (s), 1458 (m), 1400 (m), 1339 (w), 1298 (m), 1244 (m), 1190 (m), 1134 (s), 1063 (s), 1007 (s), 995 (s), 951 (m), 887 (m), 858 (w), 781 (m), 710 (m), 673 (m), 642 (m). | | | |

(1) M. Campitiello, A. Cremonini, M. A. Squillaci, S. Pieraccini, A. Ciesielski, P. Samorì, S. Masiero, “Self-Assembly of Functionalized Lipophilic Guanosines into Cation-Free Stacked Guanine-Quartets” *Journal of Organic Chemistry* 2021, 86, 9970–9978

*N*-Isobutyryl-2’-*O*-methyl-5’-*O*-(4,4’-dimethoxytrityl)-8-bromoguanosine (**3**)

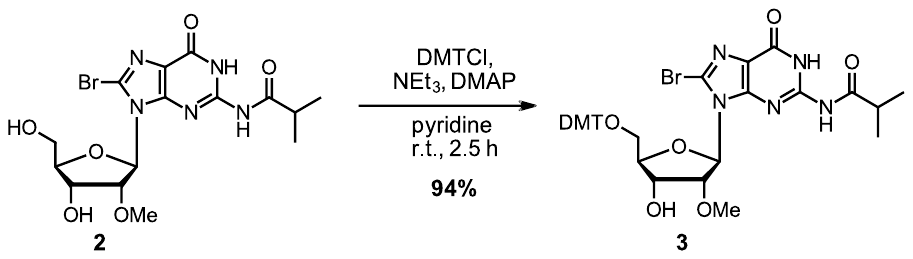

Under argon atmosphere, *N-*isobutyryl-2’-*O-*methyl-8-bromoguanosine (**2**, 2.26 g, 5.06 mmol, 1.00 equiv) was dissolved in 50 mL of abs. pyridine. NEt_3_ (719 mg, 990 μL, 7.10 mmol, 1.40 equiv), DMTCl (2.40 g, 7.09 mmol, 1.40 equiv) and DMAP (63.0 mg, 516 μmol, 0.102 equiv) were added and the reaction was stirred at rt for 2.5 h until TLC showed full conversion of the starting material. The reaction was evaporated at r.t. in the *Schlenk* vacuum, and the residue was purified via flash column chromatography on SiO_2_ (CH_2_Cl_2_/MeOH 40:1 including 1% MeOH saturated with NH_3_ 🡪 30:1 including 1% MeOH saturated with NH_3_) to obtain the product **3** in a yield of 94% (3.56 g, 4.75 mmol) as a yellow solid.

| M(C_36_H_38_BrN_5_O_8_): | 748.63 g/mol. | | 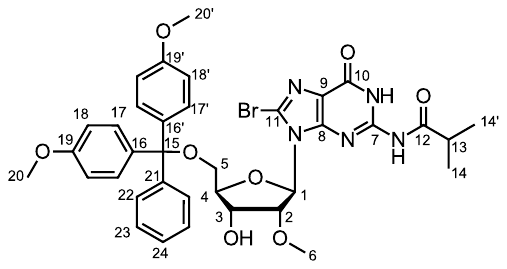 | |
| --- | --- | --- | --- | --- |
| Habitus: | Colourless solid. | |  |  |
| R_f_: | 0.48 (CH_2_Cl_2_/MeOH 20:1). | |  |  |
| ^1^H NMR: | (500 MHz, DMSO-*d*_6_) δ 12.27 – 12.11 (m, 1H, NH), 11.37 (s, 1H, NH), 7.32 – 7.28 (m, 2H, H-22), 7.20 – 7.15 (m, 7H, H-24, H-23, H-17, H-17'), 6.77 (d, *J* = 8.9 Hz, 2H, H-18), 6.72 (d, *J* = 8.9 Hz, 2H, H-18'), 5.95 – 5.93 (m, 1H, H-1), 5.19 (d, *J* = 7.2 Hz, 1H, OH), 4.73 (s, 1H, H-2), 4.54 – 4.48 (m, 1H, H-3), 4.05 (t, *J* = 6.3 Hz, 1H, H-4), 3.71 (s, 3H, H-20), 3.70 (s, 3H, H-20'), 3.45 – 3.39 (m, 1H, H-5), 3.38 (s, 3H, H-6), 3.15 (d, *J* = 10.2 Hz, 1H, H-5'), 2.73 (hept, *J* = 6.7 Hz, 1H, H-13), 1.13 (d, *J* = 6.7 Hz, 3H, H-14), 1.11 (d, *J* = 6.9 Hz, 3H, H-14'). | | | |
| ^13^C NMR: | (126 MHz, DMSO-*d*_6_) δ 180.1 (C-10), 158.0 (C-19), 157.9 (C-19'), 153.6 (C-9), 149.5 (C-8), 147.9 (C-7), 144.8 (C-21), 135.6 (C-16'), 135.5 (C-16), 129.8 (C-17), 129.6 (C-17'), 127.8 (C-22), 127.5 (C-23), 126.5 (C-24), 124.0 (C-11), 121.1 (C-9), 112.9 (C-18), 112.8 (C-18'), 88.7 (C-1), 85.3 (C-15), 84.2 (C-4), 80.1 (C-2), 69.4 (C-3), 64.1 (C-5), 58.0 (C-6), 55.0 (C-20), 54.9 (C-20'), 34.8 (C-13), 18.9 (C-14'), 18.7 (C-14). | | | |
| HR-MS (ESI): | Calc. | Exp. | | Dev. [ppm] |
|  | 748.19765 [M+H]^+^ | 748.19888 [M+H]^+^ | | +1.64 |
|  | 770.17960 [M+Na]^+^ | 770.18100 [M+Na]^+^ | | +1.82 |
| FT-IR (ATR): | 2968 (w), 2932 (w), 1682 (s), 1605 (s), 1560 (s), 1508 (s), 1458 (s), 1418 (w), 1290 (w), 1248 (s), 1177 (m), 1155 (m), 1115 (m), 1076 (m), 1034 (s), 901 (w), 826 (s), 754 (w), 725 (w), 702 (m). | | | |

*N*-Isobutyryl-2’-*O*-methyl-5’-(4,4’-dimethoxytrityl)-8-bromoguanosine-3'-[(2-cyanoethyl)-(*N*,*N*-diisopropyl)]-phosphoramidite (**4**)

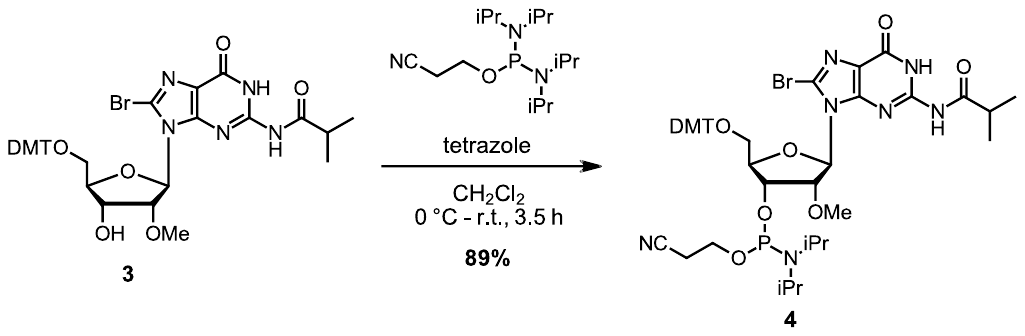

Under argon atmosphere, *N*-isobutyryl-2’-*O*-methyl-5’-*O*-(4,4’-dimethoxytrityl)-8-bromo-guanosine (**3**, 1.00 g, 1.34 mmol, 1.00 equiv) was dissolved in 12 mL of abs. CH_2_Cl_2_ and coevaporated in the Schlenk line vacuum. Then, it was redissolved in 40 mL of abs. CH_2_Cl_2_, cooled to 0 °C and 2-cyanoethylphosphor-*N*,*N*,*N*’,*N*’-tetraisopropylbisamidite (0.61 mg, 0.64 mL, 2.00 mmol, 1.50 equiv) was added. 2H-tetrazole (0.45 m in MeCN, 4.44 mL, 0.14 g, 2.03 mmol, 1.52 equiv) was added dropwise, the cooling bath was removed and the reaction was stirred at rt. After 3.5 h, the reaction mixture was washed with 50 mL of sat. NaHCO_3(aq)_. The organic layer was dried over Na_2_SO_4_ and evaporated under reduced pressure at 25 °C. The crude product was purified via flash column chromatography on SiO_2_ (cHex/acetone 2:1 + 0.7% of NH_3_-sat. MeOH) to obtain the **^Br^G_m_** phosphoramidite **4** as diastereomeric mixture in a yield of 89% (1.12 g, 1.18 mmol) as a colourless solid.

| M(C_45_H_55_BrN_7_O_9_P): | 948.85 g/mol. | | 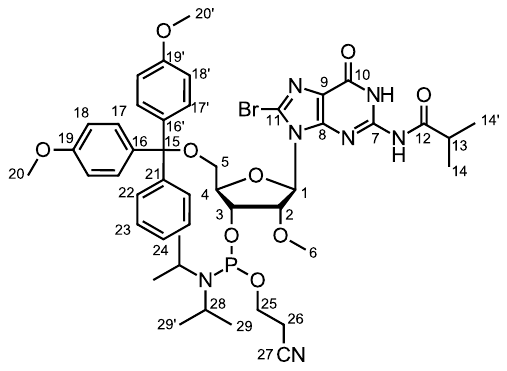 | |
| --- | --- | --- | --- | --- |
| Habitus: | Colourless solid. | |  |  |
| R_f_: | 0.40 (CH_2_Cl_2_/MeOH 30:1). | |  |  |
| ^1^H NMR: | (500 MHz, Chloroform-*d*, mixture of diastereomers) δ [ppm] 7.56 (ddd, J = 8.1, 4.2, 1.5 Hz, 2H, H-22, *dia*-H-22), 7.42 (ddd, J = 9.0, 1.8, 1.0 Hz, 4H, H-17, *dia*-H-17, H-17', *dia*-H-17'), 7.26 – 7.15 (m, 3H, H-23, *dia*-H-23, H-24, dia-H-24), 6.80 – 6.75 (m, 4H, H-18, *dia*-H-18), 6.04 – 5.99 (m, 1H, H-1, *dia*-H-1), 5.21 (dd, J = 7.8, 5.2 Hz, 0.6H, H-2), 4.94 (t, J = 6.1 Hz, 0.4H, *dia*-H-2), 4.71 – 4.64 (m, 1H, H-3, *dia*-H-3), 4.30 (t, J = 4.4 Hz, 0.4H, *dia*-H-4), 4.20 (s, 0.6H, H-4), 4.01 – 3.87 (m, 1.2H, H-25, H-25'), 3.78 – 3.74 (m, 6H, H-20, *dia*-H-20, H-20', *dia*-H-20'), 3.65 – 3.49 (m, 2.8H, H-5, *dia*-H-5, H-28, *dia*-H-28, *dia*-H-25, *dia*-H-25'), 3.48 (s, 1.8H, H-6), 3.45 (s, 1.2H, *dia*-H-6), 3.17 (dd, J = 10.7, 4.9 Hz, 0.4H, *dia*-H-5'), 3.03 (dd, J = 10.7, 3.6 Hz, 1H, 0.6H, H-5'), 2.74 – 2.65 (m, 1.2H, H-26, H-26'), 2.31 – 2.22 (m, 0.8H, *dia*-H-26, *dia*-H-26'), 1.51 – 1.47 (m, 0.4H, *dia*-H-13), 1.22 (d, J = 6.8 Hz, 2.4H, *dia*-H-29), 1.20 – 1.18 (m, 0.6H, H-13), 1.16 (dd, J = 6.8, 1.6 Hz, 6H, H-29, *dia*-H-29'), 0.97 (d, J = 6.8 Hz, 3.6H, H-29'), 0.87 (d, J = 6.8 Hz, 1.2H, *dia*-H-14), 0.81 (d, J = 6.8 Hz, 1.8H, H-14), 0.63 (d, J = 6.9 Hz, 1.2H, *dia*-H-14'), 0.52 (d, J = 6.8 Hz, 1.8H, H-14'). | | | |
| ^13^C NMR: | (126 MHz, Chloroform-*d*, mixture of diastereomers) δ [ppm] 178.9 (*dia*-C-12), 178.7 (C-12), 158.96 (C-19 or C-19'), 158.94 (C-19 or C-19'), 158.85 (*dia*-C-19 or *dia*-C-19'), 158.84 (*dia*-C-19 or *dia*-C-19'), 154.39 – 154.34 (m, C-10, *dia*-C-10), 149.7 (C-8), 149.4 (*dia*-C-8), 147.33 (*dia*-C-7), 147.27 (C-7), 145.3 (*dia*-C-21), 145.2 (C-21), 136.5 (*dia*-C-16 or *dia*-C-16'), 136.4 (C-16 or C-16'), 136.1 (*dia*-C-16 or *dia*-C-16'), 135.9 (C-16 or C-16'), 130.2 – 130.1 (m, *dia*-C-17, *dia*-C-17'), 130.1 – 130.0 (m, C-17, C-17'), 128.4 – 128.0 (m, C-23, *dia*-C-23, C-22, *dia*-C-22), 127.4 (C-24), 127.2 (*dia*-C-24), 125.37 – 125.24 (m, C-11, *dia*-C-11), 122.8 (*dia*-C-9), 122.7 (C-9), 118.0 (C-27), 117.6 (*dia*-C-27), 113.43 – 113.36 (m, C-18, C-18'), 113.27 (*dia*-C-18 or *dia*-C-18'), 113.26 (*dia*-C-18 or *dia*-C-18'), 89.0 (*dia*-C-1), 87.6 (C-1), 86.19 (C-15), 86.15 (*dia*-C-15), 84.38 (C-4), 84.35 (*dia*-C-4), 80.2 (d, J = 4.9 Hz, *dia*-C-2), 79.9 (d, J = 3.0 Hz, C-2), 71.5 (d, J = 14.5 Hz, *dia*-C-3), 69.6 (d, J = 18.2 Hz, C-3), 64.0 (*dia*-C-5), 63.5 (C-5), 59.1 (d, J = 5.4 Hz, C-6), 59.0 (d, J = 15.6 Hz, C-25), 58.6 (d, J = 3.3 Hz, *dia*-C-6), 57.04 (d, J = 19.2 Hz, *dia*-C-25), 55.46 – 55.36 (m, C-20, *dia*-C-20, C-20', *dia*-C-20'), 43.5 (d, J = 12.6 Hz, *dia*-C-28), 43.2 (d, J = 12.7 Hz, C-28), 35.9 (C-13), 35.8 (*dia*-C-13), 24.8 – 24.5 (m, C-29, *dia*-C-29, C-29', *dia*-C-29'), 20.6 (d, J = 5.2 Hz, C-26), 20.4 (d, J = 6.9 Hz, *dia*-C-26), 18.7 (*dia*-C-14), 18.6 (C-14), 18.5 (*dia*-C-14'), 18.4 (C-14'). | | | |
| ^31^P NMR: | (202 MHz, Chloroform-*d*, mixture of diastereomers) δ 150.1 (*dia*-P), 149.8 (P). | | | |
| HR-MS (ESI): | Calc. | Exp. | | Dev. [ppm] |
|  | 970.28745 [M+Na]^+^ | 970.28378 [M+Na]^+^ | | −3.78 |
| FT-IR (ATR): | 2967 (w), 2932 (w), 2874 (w), 2837 (w), 1678 (s), 1605 (m), 1557 (m), 1508 (m), 1458 (m), 1396 (w), 1364 (w), 1348 (w), 1292 (m), 1248 (s), 1223 (m), 1177 (s), 1153 (m), 1125 (m), 1076 (s), 1032 (s), 978 (s), 949 (m), 916 (m), 901 (m), 878 (m), 827 (s), 791 (s), 781 (s), 754 (m), 723 (m), 702 (s), 652 (m), 638 (m). | | | |

***LC-ESI-MS data of in-house synthesised oligonucleotides***

For LC-ESI-MS analysis an amaZon SL mass spectrometer (Bruker Daltonics) in combination with an Elute SP HPLC system (Bruker Daltonics) was used. The analysis of oligonucleotides was performed using either a Zorbax column (2.1 × 50 mm, 5 μm, SB-C18, Agilent) or an *XBridge* column (2.1 × 50 mm, 5 μm, BEH C18,Waters) was used in combination with a 10 mM triethylamine/100 mM hexafluoroisopropanol buffer as solvent A and acetonitrile as solvent B. When using the *Zorbax* column, the gradient started with isocratic 3% B for 1 min and then proceeded from 3% to 20% B in 7 min (flow rate: 0.4 ml min^-1^). Using the *XBridge* column, the gradient started with isocratic 3% B for 1 min and then proceeded from 3% to 40% B in 4 min (flow rate: 0.4 ml min^-1^).

| **Z-DNA** | 5'-C ^Br^G C ^Br^G C ^Br^G-3’ | |
| --- | --- | --- |
| 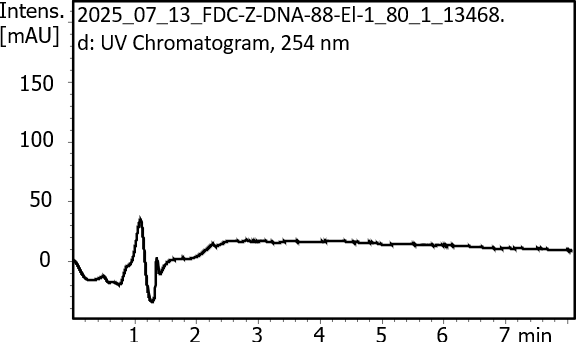 | | Column: Zorbax  M_calc._ = 1951 g/mol  M_exp._ = 1949 g/mol, 1951 g/mol |
| 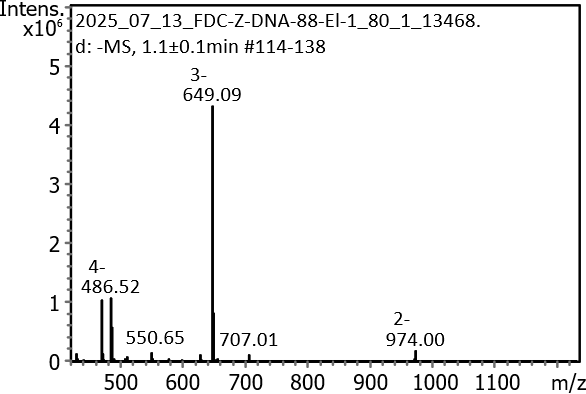 | | 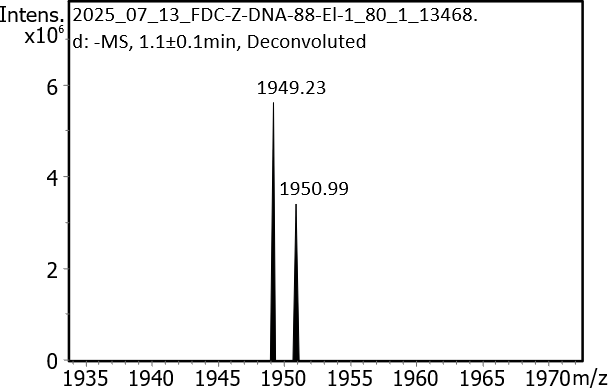 |

| **Z-RNA ^Br^G_m_** | 5'-C ^Br^G_m_ C ^Br^ G_m_ C ^Br^G_m_-3’ | |
| --- | --- | --- |
| 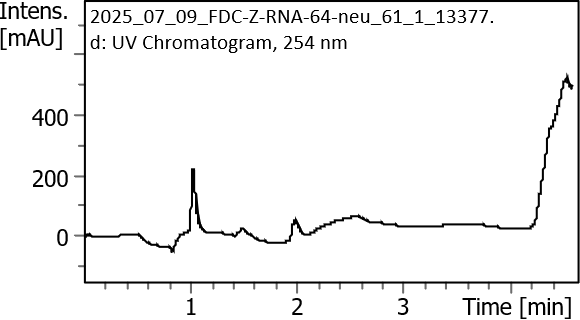 | | Column: XBridge  M_calc._ = 2075 g/mol  M_exp._ = 2073 g/mol |
| 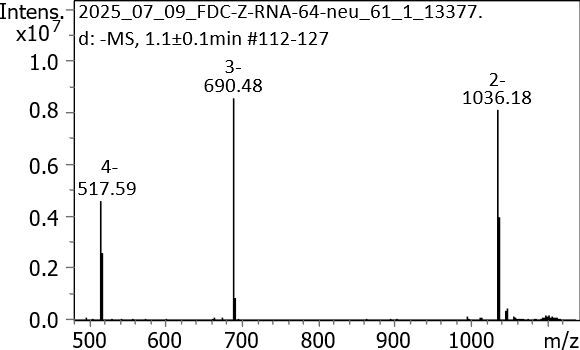 | | 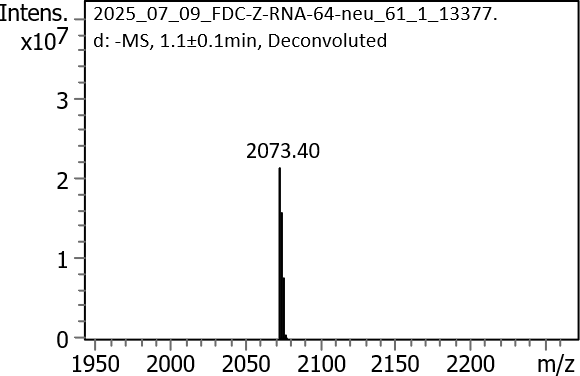 |

***Circular Dichroism and Fluorescence analysis of synthetic oligonucleotides***

B-DNA and A-RNA show opposite-sign Cotton effects in CD, with the characteristic positive band of B-DNA around 280-290 nm replaced by a negative band in A-DNA and vice versa around 250-270 nm (Figure S1A+D). The presence of negative band at 300 nm and the absence of negative band at 250 nm indicates strong presence of Z-DNA in 5’-C ^Br^G C ^Br^G C G-3’ (Figure S1B). The presence of positive bands at 250 and 280 nm indicates strong presence of Z-RNA in 5’-rC BrGm rC BrGm rC rG-3’ (Figure S1E). The strong positive bands at 260 nm in 5’-G A G G G T G G G T A G G G T G G G and 5’-rG rA rG rG rG rU rG rG rG rU rA rG rG rG rT rG rG rG indicate formation of G4 (Figure S1C+F). These structures were used to measure *in vitro* fluorescence of the DNA-binding dyes (Figure S2). The resulting fluorescence values (averages) are summarized in Table S1.

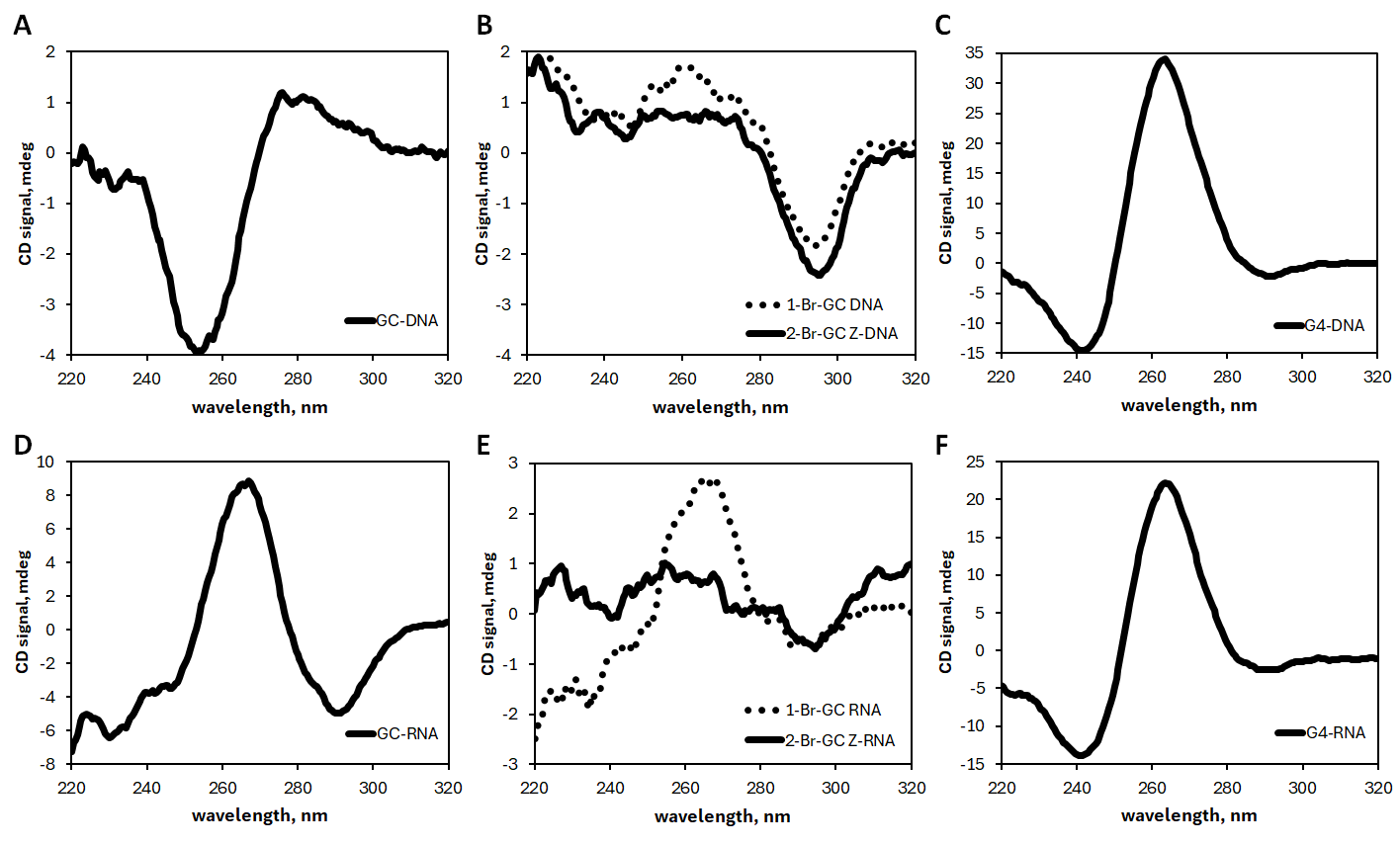

***Figure S1*** **|** **CD spectra**

Far-UV CD spectra of different nucleic acid samples. (A) B-DNA, (B) brominated DNA and Z-DNA, (C) G4-DNA, (D) A-RNA, (E) brominated RNA and Z-RNA and (F) G4-RNA. All data in mdeg.

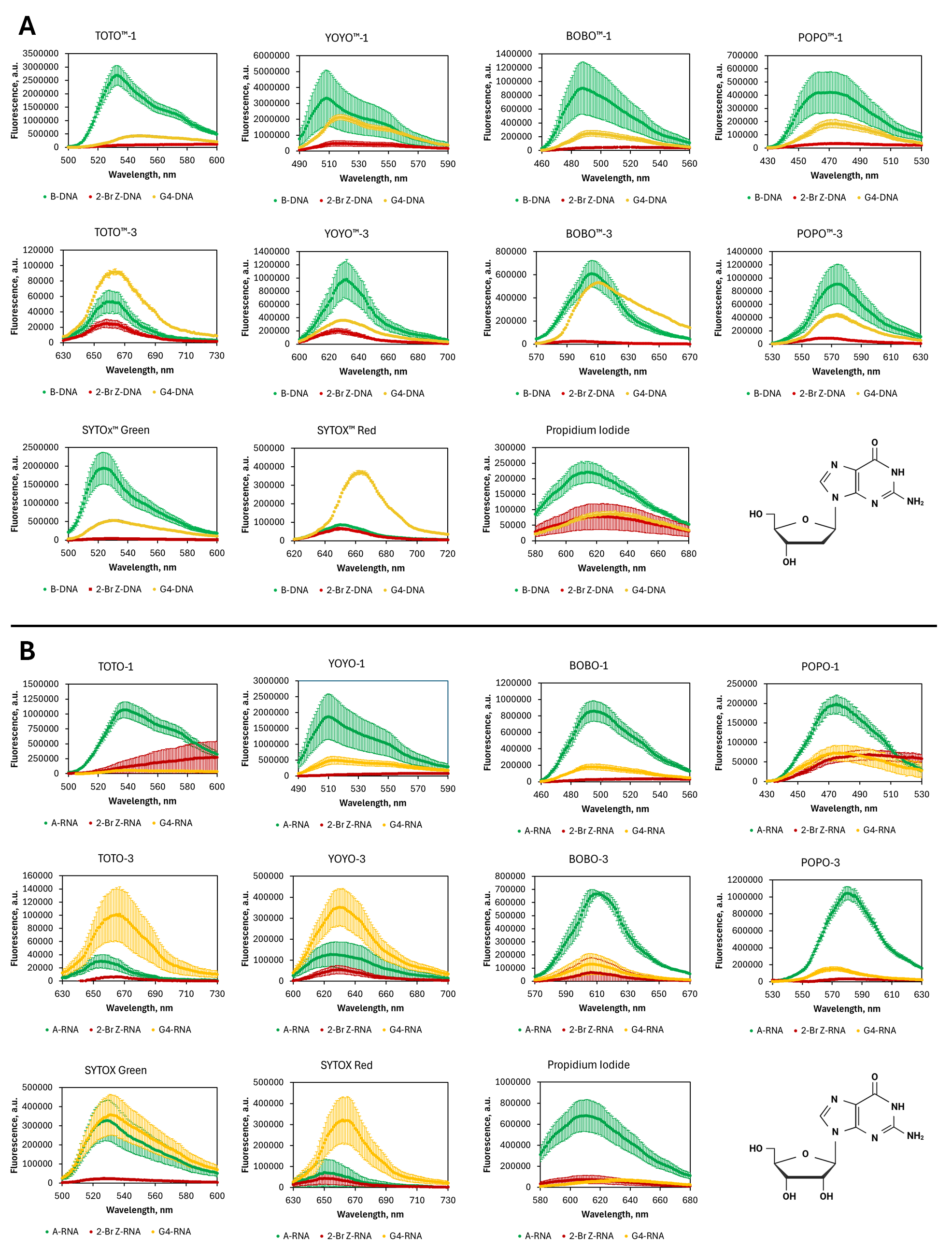

***Figure S2*** **|** **Fluorescent emission spectra**

Fluorescent emission spectra (n = 3) of NA-binding dyes in synthetic (A) DNA and (B) RNA oligos, used for constructing charts in Figure 2. Chemical structures of (A) deoxyribo- and (B) ribonucleic guanosine.

***Table S1*** **|** **Peak fluorescence values obtained from spectra in Figure S2. The values represent the average fluorescence at the peak fluorescence +/-50 nm.**

Average fluorescence values obtained from spectra in Figure S2 were used to construct Figure 2. *Indicates fluorescence in an equimolar mixture of TOTO™-1 and TOTO™-3.

|  | **B-DNA** | **Z-DNA** | **G4-DNA** | **A-RNA** | **Z-RNA** | **G4-RNA** |
| --- | --- | --- | --- | --- | --- | --- |
| **TOTO™-1** | 1.324.972 | 48.464 | 298.857 | 646.006 | 171.865 | 36.899 |
| ***TOTO™-1** | 708.089 | 15.723 | 156.555 | 325.189 | 5.628 | 80.052 |
| **TOTO™-3** | 23.561 | 8.802 | 40.661 | 11.372 | 1.901 | 47.436 |
| ***TOTO™-3** | 4.793 | 7.822 | 16.864 | 4.165 | 5.702 | 9.528 |
| **YOYO™-1** | 1.823.457 | 309.367 | 1.184.070 | 1.025.122 | 47.476 | 331.052 |
| **YOYO™-3** | 460.640 | 77.577 | 170.202 | 70.059 | 22.940 | 170.139 |
| **BOBO™-1** | 506.158 | 27.617 | 140.931 | 463.842 | 25.318 | 108.201 |
| **BOBO™-3** | 266.590 | -4.071 | 78.828 | 301.095 | 25.694 | 56.593 |
| **POPO™-1** | 253.942 | 11.555 | 101.247 | 102.653 | 46.652 | 45.519 |
| **POPO™-3** | 446.247 | 16.696 | 196.578 | 464.898 | 650 | 67.172 |
| **SYTOX™ Green** | 1.008.153 | 22.363 | 294.926 | 171.665 | 13.242 | 202.391 |
| **SYTOX™ Red** | 19.865 | 3.925 | 139.479 | 32.079 | 17.432 | 122.575 |
| **Propidium Iodide** | 130.362 | 35.662 | 59.829 | 442.063 | 48.520 | 41.988 |

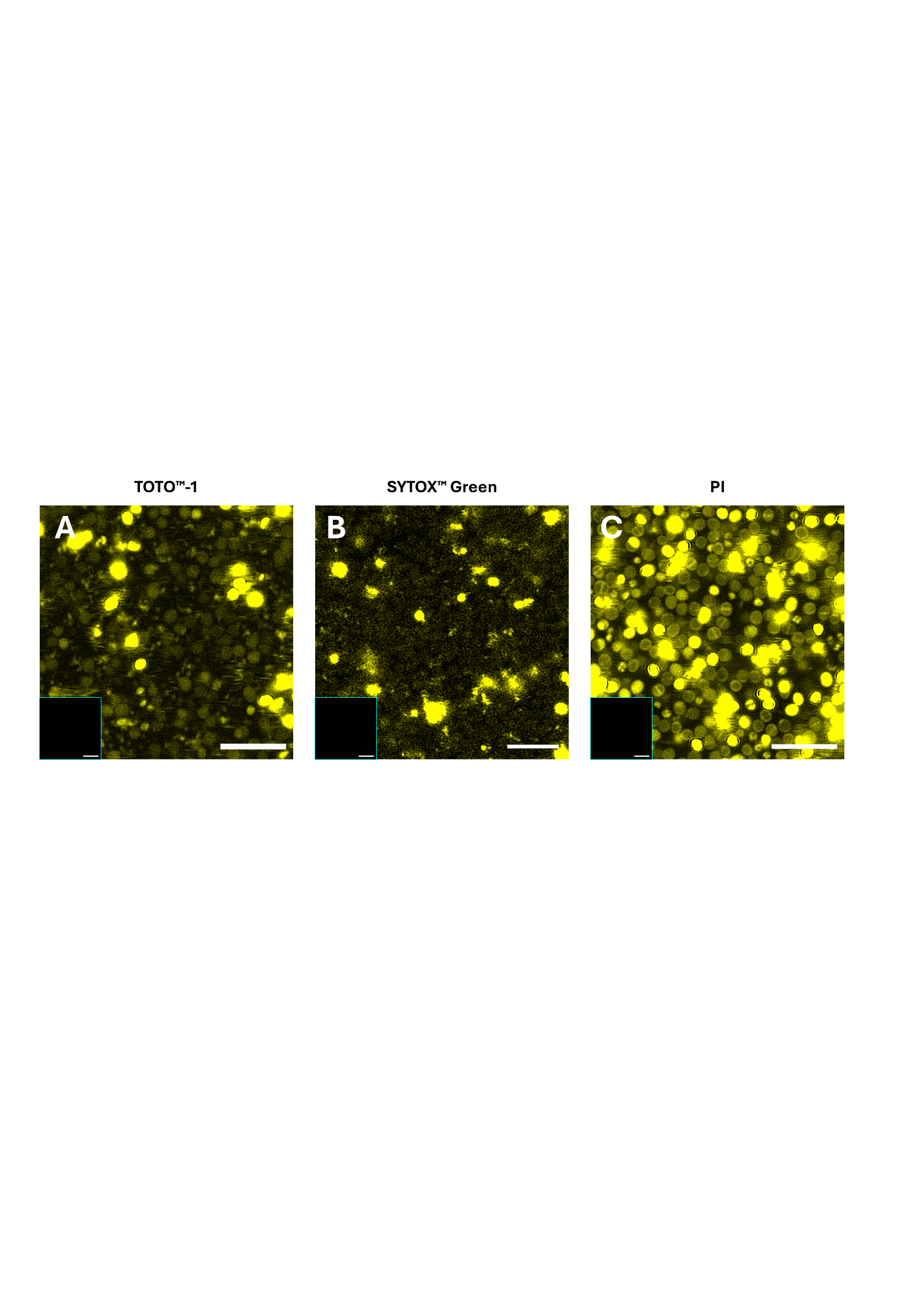

***Figure S3* | Images from Figure 4 with increased histograms.**

CLSM images from Figure 4 with increased histograms of 2-day *S. aureus* biofilms. **A** TOTO™-1 (Yellow), **B** SYTOX™ Green (Yellow), **C** PI (Yellow), **A-C** unstained control in lower left corner. Scale bars = 5 µm.

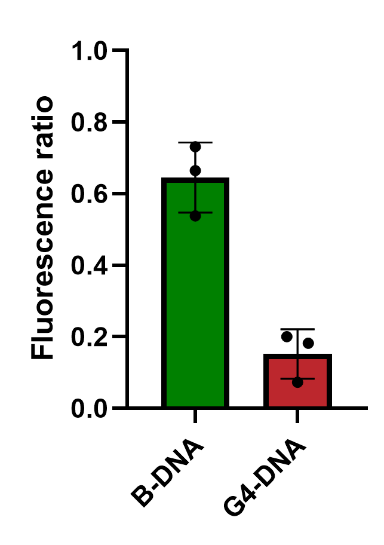

***Figure S4* | FM 4-64 reduces the signal from YOYO**™**-1 when bound to G4-DNA**

Relative fluorescence ratio between the fluorescence intensity of B-DNA (1 µM) and G4-DNA (1 µM) (Table 1) stained with YOYO™-1 (1 µM) alone and B-DNA (1 µM) and G4-DNA (1 µM) stained with YOYO™-1 (1 µM) in combination with FM 4-64 (20 µg/mL).

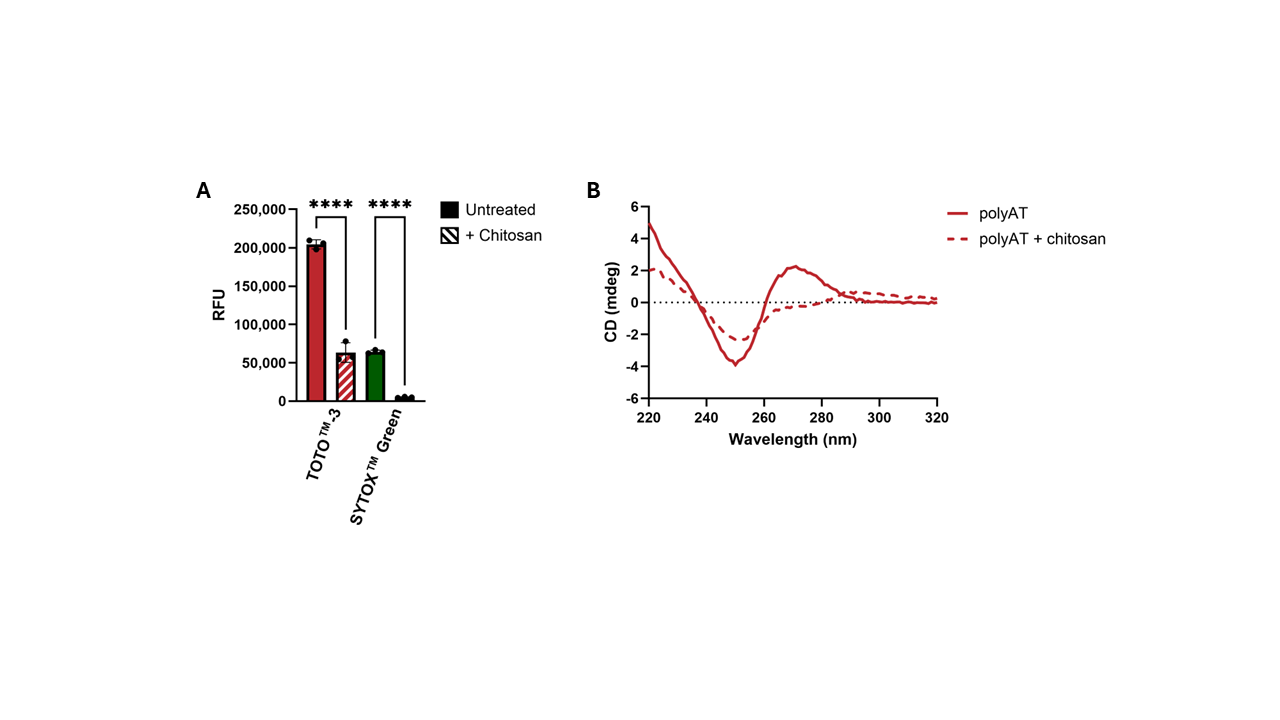

***Figure S5* | Chitosan reduces the relative fluorescence intensity of TOTO**™**-3 and SYTOX**™ **Green**

**A** Relative fluorescence intensity of TOTO™-3 and SYTOX™ Green of polyAT (5'- ATA TAT ATA TAT ATA TAT ATA TAT ATA-3') with or without chitosan. Solid bars = without chitosan, striped bars = with chitosan. **B** CD spectra of polyAT DNA (5 µM) formed with and without chitosan show B-DNA spectrum for both conditions and act as a structurally neutral control. Solid line = polyAT annealed without chitosan, dashed line = polyAT annealed with chitosan. The graph depicts ellipticity (mdeg) measured at 220-320 nm. The black dotted line indicates zero ellipticity.
